# Multicellular Organoids of the Neurovascular Blood-Brain Barrier: A New Platform for Precision Neuronanomedicine

**DOI:** 10.1101/2020.08.14.249326

**Authors:** Murali Kumarasamy, Alejandro Sosnik

**Affiliations:** Laboratory of Pharmaceutical Nanomaterials Science, Department of Materials Science and Engineering, Technion-Israel Institute of Technology, Technion City, Haifa, Israel

**Author notes:** Corresponding author: Prof. Alejandro Sosnik, Laboratory of Pharmaceutical Nanomaterials Science, Department of Materials Science and Engineering, Technion-Israel Institute of Technology, Technion City, 3200003 Haifa, Israel.

**Keywords:** Neurovascular blood-brain barrier, 3D organoids, transcriptomics, precision neuronanomedicine, nanoparticle transport, nanomedicine, nanotoxicology

## Abstract

The treatment of neurological disorders (NDs) is challenged by low drug permeability from the systemic circulation into the central nervous system (CNS) owing to the presence of the blood-brain barrier (BBB). Neuronanomedicine investigates nanotechnology strategies to target the brain and improve the therapeutic outcome in NDs. Two-dimensional adherent cell BBB models show substantial phenogenomic heterogeneity and their ability to predict the permeability of molecules and nanoparticles into the brain is extremely limited. Thus, the high-throughput screening of CNS nanomedicines relies on the use of animal models. To address this dearth, 3D organoids that mimic the *in vivo* physiology are under development. Still, there exist concerns about the standardization and scale-up of the production process, their proper characterisation, and their industrial application. In this work, we report on a novel multicellular organoid of the neurovascular blood–brain barrier (NV-BBB) that recapitulates the regulated syncytium of human endothelial cells and the function of the human BBB. For this, an advanced organoid comprising human brain microvascular endothelial cells, brain vascular pericytes and human astrocytes combined with primary neurons and microglia isolated from neonate rats is bio-fabricated without the use of an extracellular matrix. The structure and function are fully characterized by confocal laser scanning fluorescence microscopy, light sheet fluorescence microscopy, scanning transmission electron microscopy, cryogenic-scanning electron microscopy, western blotting, RNA-sequencing and quantitative gene expression by quantitative polymerase chain reaction analysis. This bulk of these self-assembloids is comprised of neural cells and microglia and the surface covered by endothelial cells that act as a biological barrier that resembles the BBB endothelium. In addition, the formation of neuron-microglia morphofunctional communication sites is confirmed. Analysis of key transcriptomic expressions show the up-regulation of selected BBB-related genes including tight junction proteins, solute carriers, transporters of the ATP-binding cassette superfamily, metabolic enzymes, and prominent basement membrane signatures. Results confirmed the more efficient cell-cell communication in 3D organoids made of multiple neural-tissue cells than in 2D endothelial cell monocultures. These multicellular organoids are utilized to screen the permeability of different polymeric, metallic, and ceramic nanoparticles. Results reveal penetration through different mechanisms such as clathrin-mediated endocytosis and distribution patterns in the organoid that depend on the nanoparticle type, highlighting the promise of this simple, reproducible and scalable multicellular NV-BBB organoid platform to investigate the BBB permeability of different nanomaterials in nanomedicine, nanosafety, and nanotoxicology.

## 1. Introduction

Neurological disorders (NDs) cause approximately 17% of the deaths worldwide and an enormous economical and societal burden.^[1,2]^ A major limitation in the treatment of NDs is that most drugs do not cross the blood-brain barrier (BBB).^[3]^ The BBB is formed by tightly bound endothelial cells and is an essential part of the neurovascular unit (NVU), a complex anatomical and functional multicellular structure comprised of a basal lamina covered with pericytes, smooth muscle cells, neurons, glia cells, an extracellular matrix (ECM), as well as a number of different neural stem/progenitor cells.^[4]^ Understanding the central nervous system (CNS) pathways in health and disease as well as the evaluation of novel neurotherapeutics has been challenging due to the complexity of the NVU.^[5]^

The use of nanotechnology to improve the delivery of neurotherapeutics to the CNS, a field coined neuronanomedicine, has emerged as one of the most dynamic research areas in nanomedicine.^[6]^ Different strategies have been investigated to surpass the BBB by systemic (e.g., intravenous) and local (e.g., nasal) administration routes.^[6,7]^ More recently, nanotoxicology, the discipline that investigates the toxicity of nanomaterials, has devoted efforts to develop reliable models to assess the detrimental interaction of different nanomaterials with the CNS upon intentional or unintended exposure.^[8]^ The systematic investigation of the biocompatibility, safety, permeability, and efficacy of neuronanomedicines remains mostly limited to *in vivo* experiments. However, the complex physiology of animal models challenges the conduction of permeability and mechanistic studies to understand the transport of nanoparticles (NPs) into the CNS.^[9]^

For years, endothelial cell monolayers cultured on semipermeable membrane well-plates (e.g., Transwell^®^) have been the most commonly and widely used *in vitro* models of the BBB.^[10]^ They utilize user-friendly setups, are scalable and enable high-throughput screening. However, they have been questioned because they cannot mimic the complex 3D cellular structure, the physiological microenvironment, the cellular phenotype and the cell-cell interactions in the NVU.^[11]^ In addition, they exhibit edge effects that hinder cell growth, especially at the plate edges. Thus, the surface area of the semipermeable membrane may not be fully covered by cells, which artificially increases the permeability. More recently, the development of 3D cell culture models has gained attention to investigate the transport of different neurotherapeutics into the brain.^[12]^ These models also represent a valuable tool to investigate pathophysiological pathways in the CNS.^[13]^ Advantages of 3D organoids include easy and reproducible culture, miniature scale, small reagent volumes, low relative cost, reproducibility, and scalability. Furthermore, they reduce animal experimentation.^[14]^ Most of these 3D models are one-cell or three-cell cultures and fail to fully recreate the NVU-BBB.^[15]^ One usually missing cell type in these models is the so-called ‘third element’ of the CNS, which is in fact resident macrophages (microglia) that constitute 10-15% of the total cells in the brain.^[16]^

We recently reported on the possible role of olfactory microglia in the nose-to-brain transport of different nanomaterials.^[17]^ This mechanism might be also exploited by pathogens such as the coronavirus-19 (COVID-19) to enter the CNS through the olfactory epithelium,^[18]^ though the key cell players in this pathway are not known yet. In addition, glial cells (e.g., astrocytes), microglia and neurons are actively involved in neuro-haemostasis and regulate the transmission of electrical signals in the brain.^[19]^ Despite recent advances in the development of cell spheroids and organoids of the BBB, many drawbacks persist. They usually lack the essential BBB cellular milieu, including microglia, six distinct cortical layers, and endothelial vasculature. Moreover, the limited formation of microglia and mature neurons limits its utility for specific *in vitro* ND models. Thus, the application of these simplified 3D models to understand how different neurotherapeutics in general and neuronanomedicines in particular are transported across the BBB and interact with phagocytic cells likely involved in their uptake and eventually their clearance remains a significant scientific challenge.^[5,12d]^ In this scenario, the development of more robust, predictive, and cost-effective *in vitro* models that recapitulate better the complex cell-cell interactions in the BBB and the NVU function is called for.

In this work, we investigate a novel multicellular organoid comprised of human brain microvascular endothelial cells (hBMECs), human brain vascular pericytes (hBVPs) and human astrocytes (hAs) combined with primary neurons and microglia isolated from neonate rats that was produced by spontaneous cell self-assembly and without the use of an extracellular matrix (ECM) such as Matrigel^®^. The structure of the organoids and the presence of cell-cell interactions is characterized through the integration of confocal laser scanning fluorescence microscopy (CLSFM), light sheet fluorescence microscopy (LSFM), scanning transmission electron microscopy (STEM) and cryogenic-scanning electron microscopy (cryo-SEM).^[20]^ In addition, the up-regulation of selected key BBB-related genes that code for tight junction proteins, solute carriers (SLCs), transporters of the ATP-binding cassette superfamily (ABCs), metabolic enzymes and proteins of the basal membrane were screened by western blotting, RNA-sequencing (RNA-Seq) and gene expression by quantitative polymerase chain reaction (qPCR) analysis. Finally, these multicellular organoids were used to characterize the permeability of different model polymeric, metallic, and ceramic NPs. Overall results support the promise of this simple and scalable multicellular NV-BBB organoid platform to investigate the interaction of nanomaterials with the BBB in neuronanomedicine and nanotoxicology.

## 2. Results and Discussion

### 2.1. Establishment and structural characterization of the neurovascular-BBB

Multicellular neurovascular organoids are one of the most promising surrogate *in vitro* models in translational neuronanomedicine, overcoming some of the shortcomings of monocellular 2D and 3D models.^[21]^ However, they do not incorporate microglia cells, which mediate immune responses in the CNS by acting as macrophages and clearing cellular debris, dead neurons, and up-taking foreign particles. In addition, they usually require complex fabrication procedures.

In previous studies, we used BBB endothelial and olfactory neuroepithelial cells isolated from adult and neonate rat to study the transport of different polymeric nanoparticles.^[17,22]^ The aim of the present work was to extend these investigations and to develop a platform of multicellular organoids as a tool to assess the interaction of different types of nanomaterials with the BBB. For this, we used a simple self-assembly method without ECM to bio-fabricate organoids that combine three human cell types, namely hCMEC/D3, hBVPs and hAs, and incorporated two primary rat cell types, neurons that form synapses and neuronal networks and microglia cells involved in the clearance of particulate matter (**Figure 1A**).

**Figure 1.**
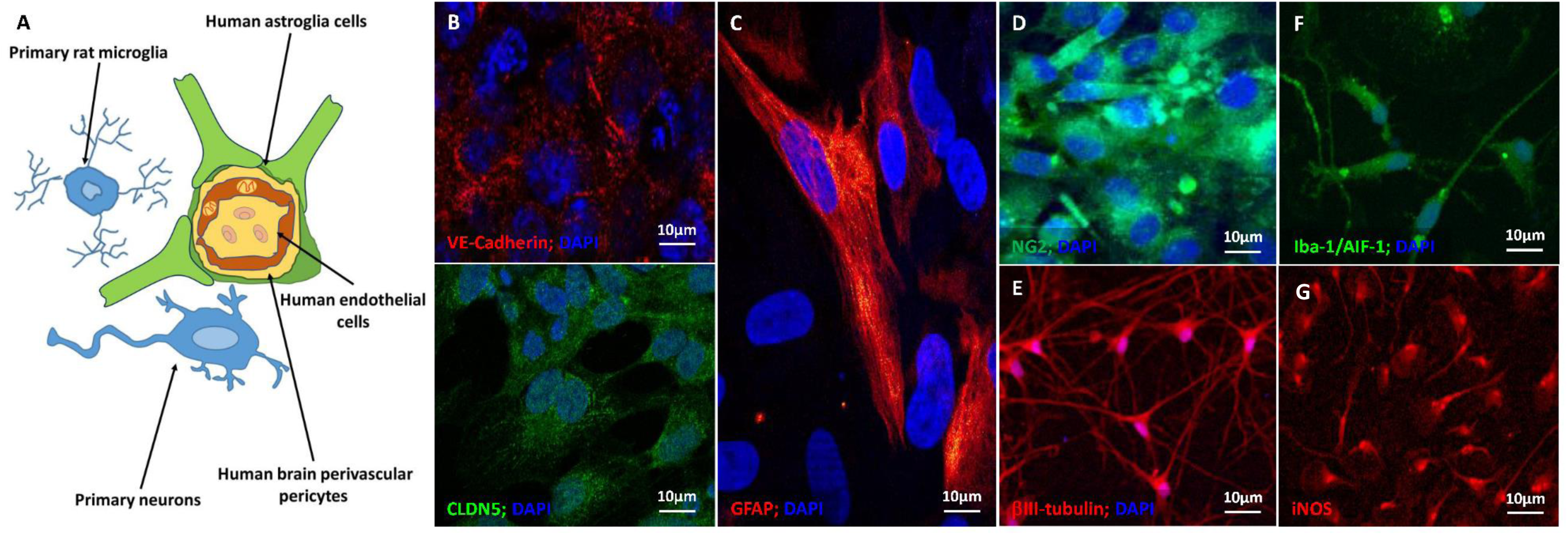
Immunocytochemical characterization of the human and rat cells used in the bio-fabrication of neurovascular organoids cultured in flat well-plates. (A) Scheme of the neurovascular units and (B-G) CLSFM micrographs of (B) hCMEC/D3 endothelial cells cultured on collagen-coated glass coverslips and their specific VE-cadherin (red) and CLDN5 (green) staining, (C) GFAP-positive primary hAs, (D) NG2-positive hBVPs (green), (E) βIII-tubulin-positive primary rat neurons (red), (F) Iba-1/AIF-1-positive primary rat microglia and (G) iNOS-positive primary rat microglia. Cell nuclei in B-F are stained with DAPI (blue).

Before organoid fabrication, we characterized the five different neural-tissue cell types by immunocytochemical staining. hCMEC/D3 cells are derived from human temporal lobe endothelial microvessels and produce two characteristic proteins of adherens and tight junctions, VE-cadherin and CLDN5, respectively (**Figure 1B**). Primary hAs express the filament protein GFAP (**Figure 1C**) and hBVPs the NG2 proteoglycan (**Figure 1D**). Primary neurons (**Figure 1E**) and microglia (**Figure 1F,G**) from neurogenic and non-neurogenic regions of neonate rat brains express βIII-tubulin, which is a microtubule element almost exclusive of neurons, and Iba-1/AIF-1 and iNOS which are overexpressed in activated microglia. Primary neurons are also positive for MAP-2 (data not shown). In these experiments, microglia changed the morphology from ramified characteristic of quiescent cells to a more flattened macrophage-like one (**Figure 1F,G**).

In addition, we investigated the interaction of rat microglia and hCMEC/D3 in 2-cell spheroids. Qualitative analysis suggests that there are not detrimental interspecies interactions^[23]^ such as bidirectional and permanent communications between them (**Supplementary Figure S1**).

After the characterization of the individual cells, we used them to bio-fabricate multicellular organoids that resemble the complex brain structure and utilized them to assess the permeability of the nanoparticles. We hypothesized that the phenotype of hCMEC/D3 in multicellular organoids will mimic better their physiology and function in the BBB. To serve as a high throughput screening tool in neuronanomedicine, the bio-fabrication process needs to be simple, cheap, reproducible, robust, and eventually scalable. We adapted a previously reported method in which all the cells are mixed and cultured on agarose gel or round bottom well-plates (see experimental section). After 2-3 days of incubation, spherical organoids formed, and they were fully characterized at day 5 (**Figure 2**). 3D cultures were small spherical bodies with radially distributed cells, whereas monocultures in flat well-plates exhibited enlarged cell bodies with less and short processes.

**Figure 2.**
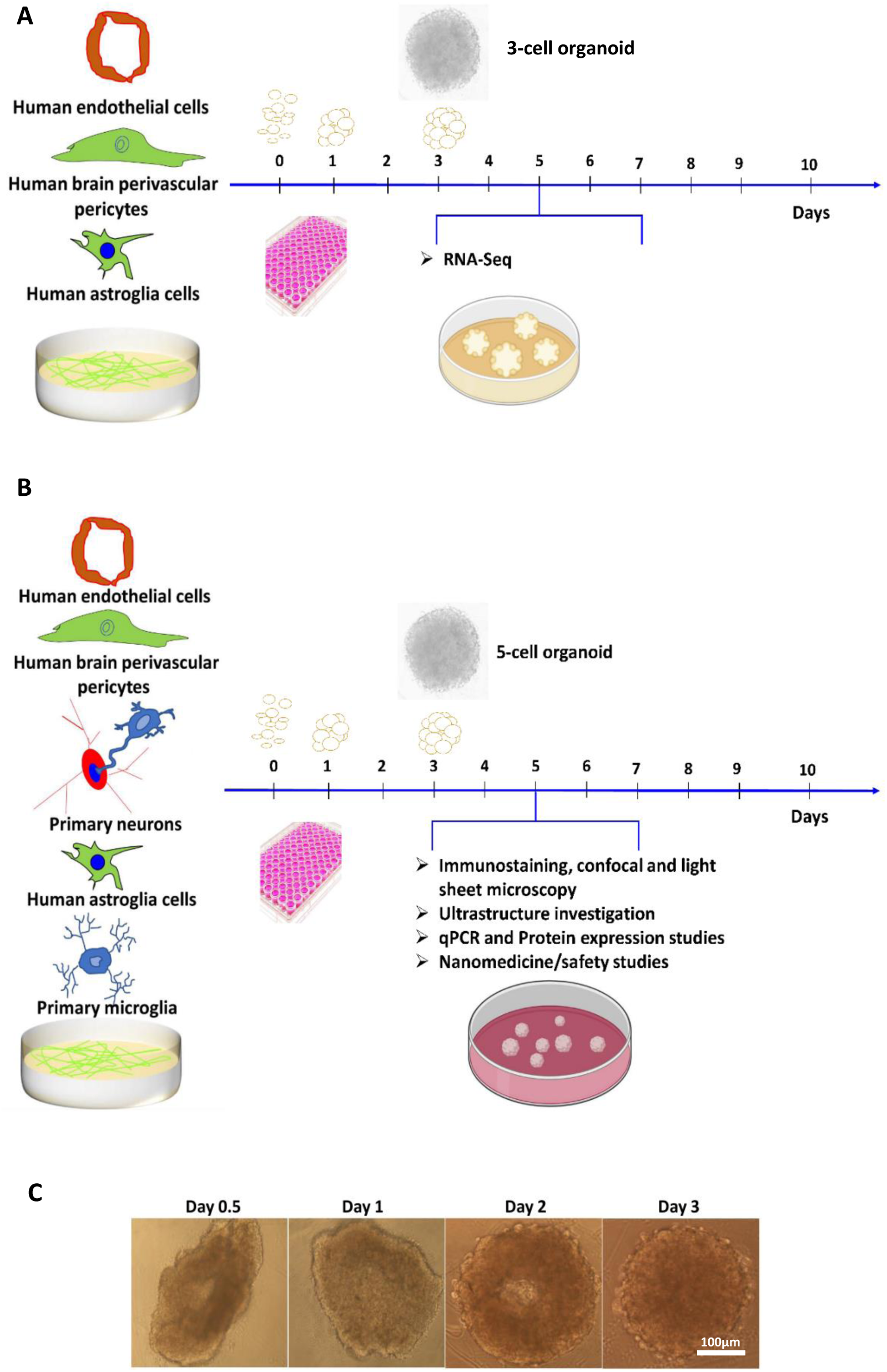
Bio-fabrication and characterization of neurovascular organoids. Scheme to produce (A) 3-Cell organoids and (B) 5-cell organoids. (C) Spherical multicellular organoids were formed within 2-3 days and fully characterized at day 5.

Upon production, organoids were immunostained to reveal the cellular architecture by CLSFM (**Figure 3A-H**). hCMEC/D3 endothelial cells together with pericytes appeared to form a surface monolayer tightly encasing the rest of the cells in the spheroid and they expressed VE-cadherin (**Figure 3A**) and CLDN5 (**Figure 3B**) which are characteristic of adherens and tight junctions, respectively. In addition, organoids showed a strong immunoreactivity for GFAP (**Figure 3C**) that is characteristic of hAs, *f*-actin of key functional proteins and the AQP4 water channel at the astrocyte end-foot covering the whole organoid (**Figure 3D**) that is crucial for the function of the BBB, the NG2 proteoglycan of mature hBVPs (**Figure 3E**), βIII-tubulin and MAP-2 of primary rat neurons (**Figure 3F,G**, respectively), and Iba-1/AIF-1 of primary rat microglia (**Figure 3H**). Microglial processes at the neuro-glia junctions could potentially monitor and protect neuronal functions.

**Figure 3.**
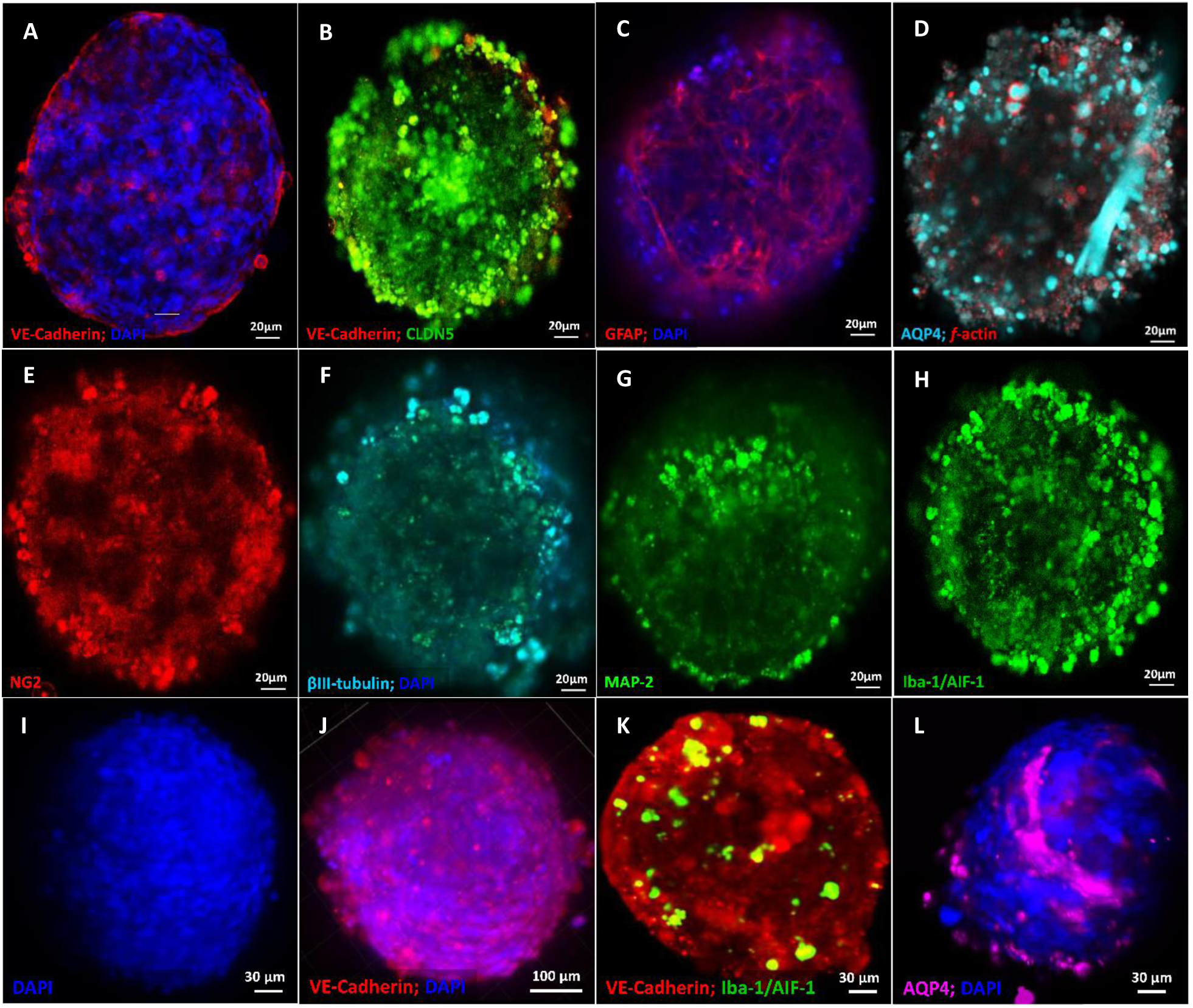
Immunocytochemical characterization of the bio-fabricated 5-cell neurovascular organoids. Representative (A-H) CLSFM and (I-L) LSFM micrographs. Organoids showed the expression of characteristic markers of (A,B,J,K) hCMEC/D3 endothelial cells, (C,D,L) primary hAs, (E) hBVPs, (F,G) primary rat neuron and (H,K) primary rat microglia. Cell nuclei in A, C, F, I, J and L were stained with DAPI (blue).

Previous experiments demonstrated that CLSFM is not the most appropriate technique to monitor the inner cellular structure and the diffusion of fluorescently-labeled nanoparticles into the organoids because it only allows to scan a Z-stack depth of 100 µm. Imaging in deeper layers is time-consuming and not feasible. The visualization of the whole multicellular organoid could be conducted more efficiently and in a short time by 3D tomography using LSFM; this method enables the detection of fluorescence signals and the imaging of the sample as deep as 1 mm, and thus of the organoid core.^[24]^ LSFM evidenced that our organoids are a solid cellular structure (**Figure 3I**) where hCMEC/D3 endothelial cells cover almost completely and uniformly the surface, forming adherens junctions that are a fundamental structure to govern the permeability into the CNS (**Figure 3J,K**). This observation was confirmed by CLSFM (**Supplementary Figure S2**). In addition, microglia cells express Iba-1/AIF-1 (**Figure 3K**), a microglia/macrophage-specific Ca^2+^-binding protein that participates in membrane ruffling and phagocytosis in activated microglia.^[25]^ The staining of AQP4 confirmed the presence of abundant filamentous bundles characteristic of hAs in the spheroid core (**Figure 3L**). Perivascular regions are also essential to mimic the physiology and function of the BBB along with the vascular endothelial barrier and astrocytic end-feet. For instance, astrocytes restore their phenotype in a 3D culture.^[26]^ In line with previous works, we confirmed that hAs cultured in multicellular organoids exhibit a ramified phenotype that resembles cortical astrocytic networks, as opposed to 2D cultures where this cell type exhibited enlarged cell bodies, with less and shorter processes (**Figure 1C**). This phenotype may contribute to regulate the permeability of the BBB, either independently or in concert with other neighbouring neuron–glia cells.^[27]^ Furthermore, we performed Western blot analysis and confirmed unequivocally the expression of key proteins such as VE-cadherin (120 kD), β-III tubulin (55 kD), GFAP (52 kD) and Iba-1/AIF-1 (19 kD) (**Supplementary Figure S3**). To ensure that the level of total protein seeding in all the runs was similar, we stained the gels with Coomassie blue staining directly after running gel electrophoresis (data not shown).

### 2.2. Cellular organization and ultrastructure

To gain a deeper understanding on the organoid ultrastructure and fundamental cell-cell interactions, we analyzed selected organoid samples by STEM, and characterized endothelial, neuronal, glia and pericyte compartments, the formation of synapses, morphofunctional communication sites between microglial processes, and neuronal cell bodies and the recruitment of phagocytic cells. This technique is advantageous because immunocytochemical staining is not needed. In addition, it can be utilized to visualize the interaction of the different cells with metallic NPs (see below).

Consistent with our observations using CLSFM and LSFM, hCMEC/D3 endothelial cells are characterized by an elongated shape on the organoid surface, irregular nuclei, and tightly regulated syncytium of the outer organoid surface (**Figure 4A**). A detailed ultrastructural analysis at the point of cell–cell connections between endothelial cells on the surface of each organoid revealed the formation of dense strands of tight and adherens junctions (**Figure 4B,C**). The outer endothelial cell-covered surface would recapitulate the phenotype of these cells in the BBB and could govern the transport of NPs and other nano-objects (e.g., viruses) into the organoid, and serve as a tool to compare their permeability.

**Figure 4.**
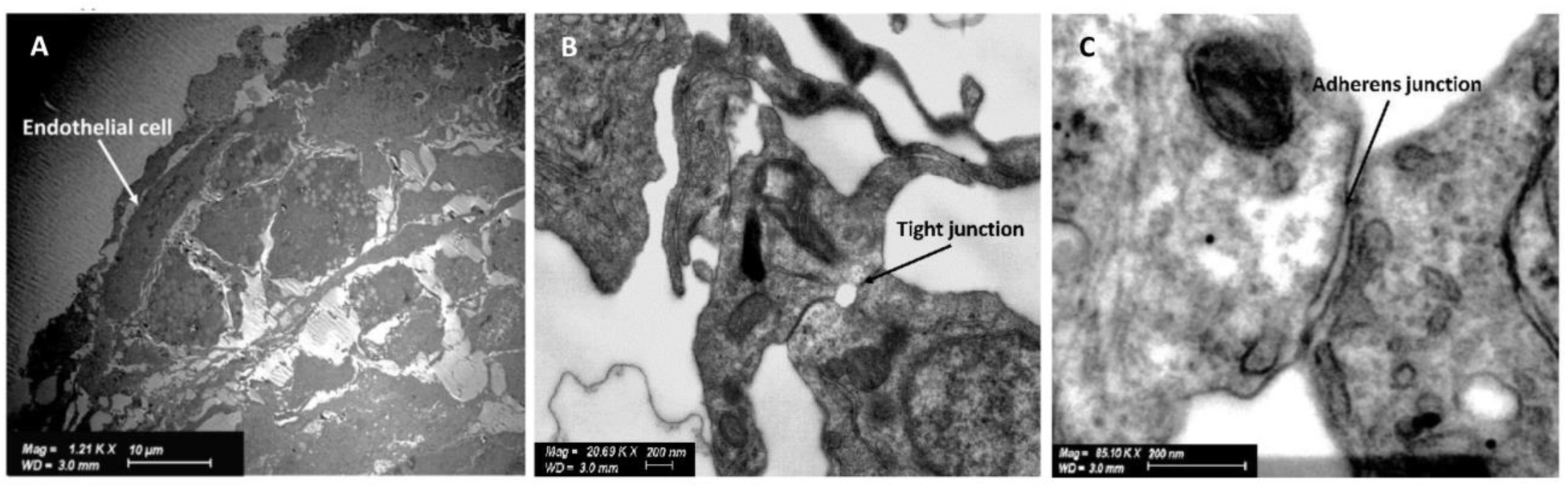
Ultrastructural characterization of hCMEC/D3 endothelial cells in 5-cell neurovascular organoids. Representative STEM micrographs showing (A) the organization of hCMEC/D3 endothelial cells on the spheroid surface, and the formation of (B) tight junctions and (C) adherens junctions.

The brain is comprised of billions of biological wires that communicate with each other through an intricate web of axons and dendrites. We observed the presence of this and other organelles (e.g., Golgi canals) in primary neurons (**Figure 5A**). Neurons may also sense and respond to their environment via a specialized organelle called primary cilium, which is composed of nine microtubule pairs on most neurons and a unique solitary, microtubule-based appendage linked to neuronal signalling. The primary cilium is thought to act as an antenna surveying the extracellular milieu, accepting, and transmitting various signals to the neighbouring cells (**Figure 5B**). In addition, in ultrastructure cross-sections, myelinated axons exhibited nearly circular profiles surrounded by a spirally wound multilamellar electrical insulator, which enables the electrical impulses between these biological wires to travel back and forth quickly (**Figure 5C**). Moreover, this analysis provided an experimental proof that microglia, a CNS macrophage, interacts with primary neurons and their synapses (**Figure 5D**).

**Figure 5.**
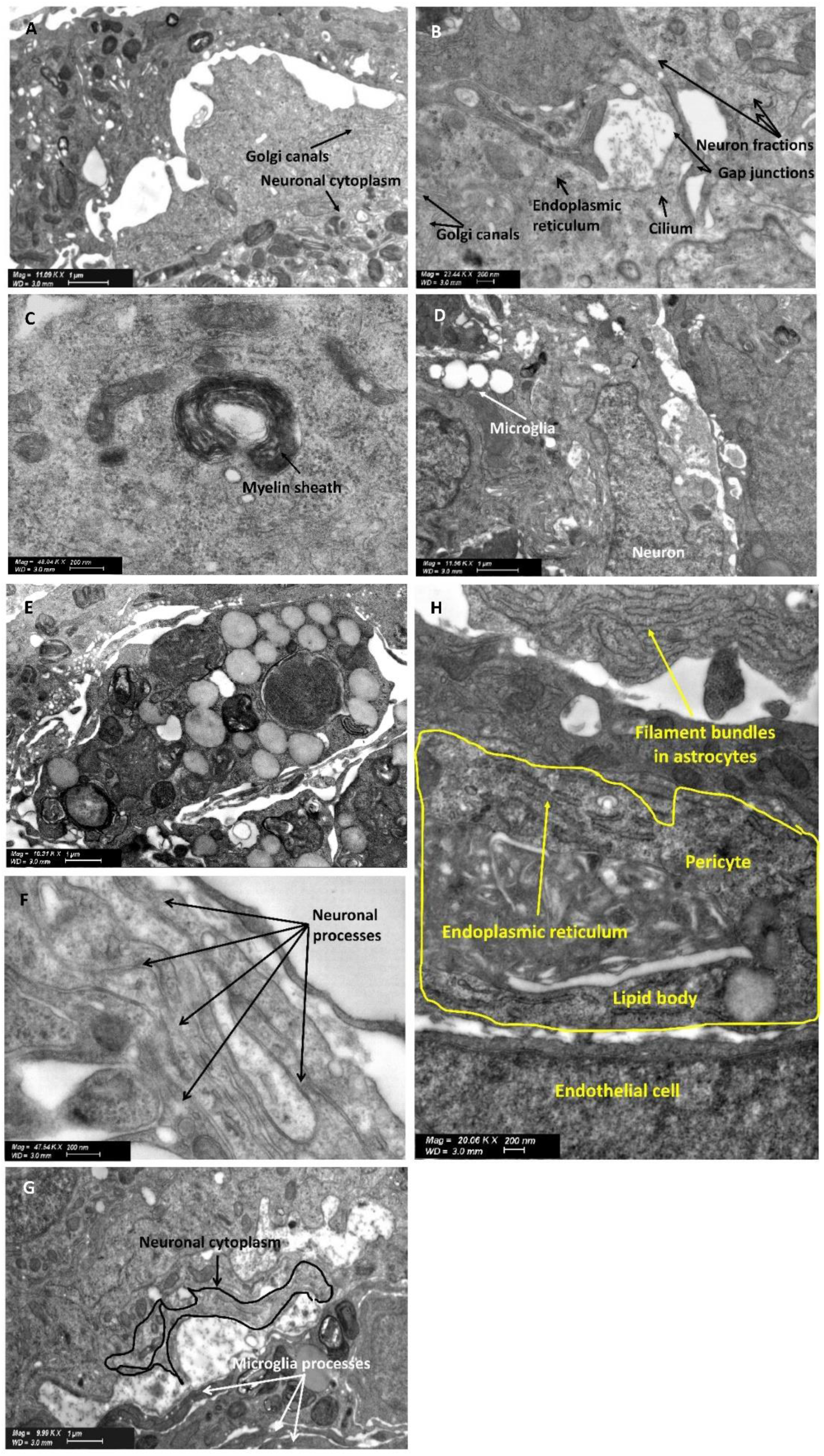
Ultrastructural characterization of neurons and microglia in 5-cell neurovascular organoids. Representative STEM micrographs showing (A) Part of a neuronal cytoplasm and the presence of Golgi canals, (B) neuronal fractions, the primary neural specific cilium lined on the surface of the organoid, Golgi canals, endoplasmic reticulum and ribosomes, (C) a myelinated sheath in the organoids along with electron dense Nissl bodies of the neuronal cytoplasm (indicated with dotted circles), (D) microglia and neuronal junctions, (E) lipid bodies characteristic of microglia, (F) neuronal processes, (G) microglial processes connecting specialized areas of the neuronal cytoplasm, and (H) endothelial cell process extending to form a junction with an overlying pericyte.

Furthermore, this technique provided a direct ultrastructural evidence that neurons are essential for immune cell-neuron communication (**Figure 5D**), which is in line the neuroprotective effect of microglia. Microglial cell bodies can be discerned from other cell types by a smaller size (3–6 μm), electron-dense cytoplasm, bean-shaped nuclei and the accumulation of light inclusions known as lipid bodies (**Figure 5D,E**). They also display a ring of cytoplasm separating the nucleus from the cell membrane, contain few organelles within a single ultrathin section, and a distinct thick, dark band of electron-dense heterochromatin located near the nuclear envelope with pockets of compact heterochromatin nets throughout the nucleus (**Figure 5E**). Microglia play the role of dynamic sensor of the brain environment by forming motile processes and by constantly interacting with neighbouring neurons, promote proper neuronal wiring and activity, and protect them from external insults. Our results confirmed the presence of microglial processes, and morphofunctional microglia-neuronal communications in the organoids (**Figure 5F**).

A critical phase in the development of the CNS is cell migration, often over long distances, from their origin place to their mature position. Organoids displayed the presence of neuronal processes that would be consistent with neuronal migration (**Figure 5G**), an essential process for the development of the nervous system. Astrocytes, like other glial cells, have been commonly presumed as mere support for the key players in brain functioning, neurons, and the biological wires in the CNS. At the ultrastructural level, astrocytes can be identified by an irregular, stellate shape, with numerous glycogen granules, bundles of intermediate filaments, and a relatively clear cytoplasm (**Figure 5H**). STEM studies confirmed the formation of pericyte-endothelial cell connections that have a peg and socket arrangement (**Figure 5H**) and that enables signal transmission mediated by the release of VE-cadherin (**Figure 3A,B,J,K**). Finally, we visualized the 3-cell BBB organoids with cryo-SEM to study the ultrastructural morphometry at near-native context. Cryo-tomograms showed filament bundles between the cells (**Supplementary Figure S4**).

### 2.3. RNA-sequencing

One of the challenges in the production of multicellular NVU organoids is to achieve an endothelial cell phenotype that resembles the function *in vivo* because the BBB endothelium regulates the transport of soluble and particulate matter into the CNS. We anticipated that 3D co-culture with hBVPs and hAs would result in a more physiological endothelial cell phenotype.

To analyze whether our multicellular organoids exhibit physiological characteristics of the *in vivo* BBB and constitute a functional barrier or not, we evaluated and compared transcriptome expression by RNA-Seq at day 5. Owing to interspecies variabilities,^[28]^ for these studies, we used 3-cell organoids comprised on only human cells, namely hCMEC/D3 endothelial cells, hAs and hBVPs, and compared them to 2D and 3D endothelial cell monocultures; endothelial cell monolayers are the most common *in vitro* model of BBB. The quality of the extracted RNA was assessed by 1% agarose gel electrophoresis and the quantity and the purity (quality control) of the RNA samples by using the Qubit^®^ and TapeStation; all the samples showed RNA integrity (RIN) values above 8 (**Supplementary Table S1**).

To visualize transcriptomic differences between 3-human cell BBB organoids, endothelial cell (3D) organoids, and endothelial flat (2D) cultures in genes coding for key proteins, a heatmap was generated showing log2 (fold-change) >2 (**Figure 6A-E**); for each group three independent biological replicates were analyzed (n = 3).

**Figure 6.**
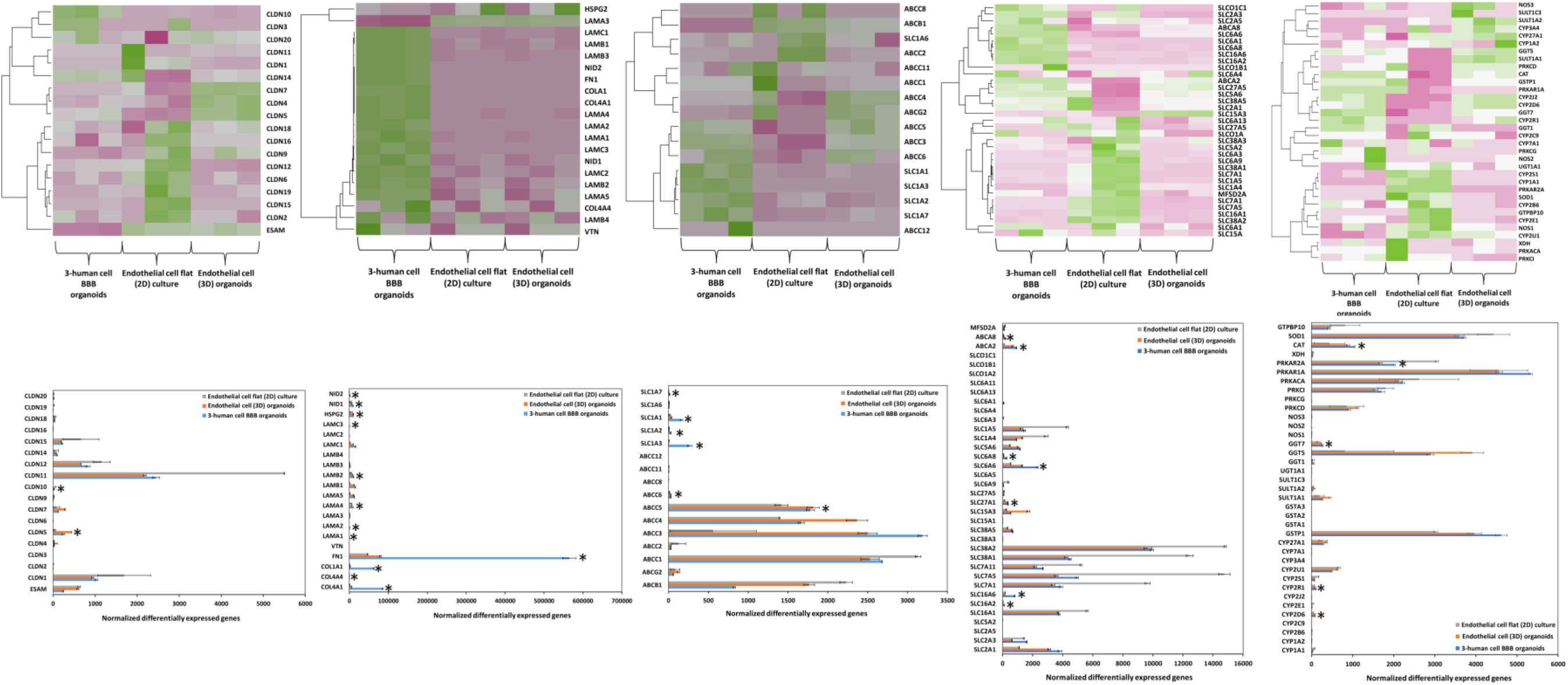
Transcriptomic (RNA-Seq) analysis. Heatmap of RNA-Seq and differentially expressed genes (DEGs) upregulated analysis of 3-human cell organoids and 2D and 3D endothelial cell monocultures; (n = 3 for each culture condition). Green and pink indicate up-regulation and down-regulation, respectively. Average of hierarchical clustering indicates the interclass correlation between all three groups. Selected differential expression of genes encoding for (A,F) tight junction proteins, (B,G) extracellular matrix (ECM) proteins, (C,D,H,I) ABC efflux transporters, solute carriers (SLC) and other nutrient transporters and (E,J) metabolic enzymes. *Significantly differentially expressed genes (DEG) (p_adj_ <0.05, | Fold Change | >2, base Mean >= 20). To provide optional filtering criteria in addition to the p_adj_, additional criteria of |Fold change| >2 (|log2 Fold Change| >1) and average expression level higher than 20 (base Mean >20) were used.

Based on the criteria given in the experimental section, the number of reads ranged from 21,842,753 to 27,419,486 per sample (**Tables 1,2** of **Supplementary experimental file**). We performed principal components analysis (PCA) on variable groups (excluding the outlier endothelial cell flat culture), to identify genes that are most informative for defining cell subpopulations (**Supplementary Figure S5**). PCA plots were useful for visualizing the overall effect of experimental covariates and effects on each model. The percentage of uniquely mapped reads ranged from 93.87 to 95.28 per sample (**Table 3** of **Supplementary experimental file**). As a first step to compare the transcriptomic effects on 3-human cell BBB organoids, endothelial cell (3D) organoids and endothelial cell (2D) monolayers, comparative data was generated to display the number of differentially expressed transcripts. To assess whether the transcript was similarly altered in the transcriptomes produced in response to cell-cell interactions including endothelial-astrocyte-pericyte ones in the 3-cell organoid, a more detailed comparison was carried out by showing a heatmap that represents the quantitative fold change value under each organoid model (**Figure 6A-E**). These initial analyses revealed that multicellular BBB organoids and both 2D and 3D endothelial cell monocultures express key genes (**Figure 6**; **Supplementary file**). To determine whether these gene expression profiles were statistically different between the three groups, we analyzed RNA-Seq data by using the Pearson correlation coefficient and unsupervised hierarchical clustering. According to heatmaps, the gene expression profile of 3-human cell organoids generally differed from that 2D and 3D endothelial cell monocultures.

The three groups showed close distance within samples. We assume that there is a different cell milieu and that in the 3-cell organoids, most transcripts stem from endothelial cells. Next, we confirmed the differentially expressed genes (DEGs) between the three different groups. We set the threshold to p_adj_ <0.05 and FC >2. Results showed that 7314 genes were up-regulated in 3-cell organoids with respect to endothelial cell flat (2D) cultures, 3966 genes were up-regulated in 3-cell organoids with respect to endothelial cell (3D) organoids, 6290 genes were up-regulated in endothelial flat (2D) cell cultures with respect to endothelial cell (3D) organoids, and 6273 genes were down-regulated in 3-cell organoids with respect to endothelial cell flat (2D) cultures (**Supplementary Table S2**). Due to the relevance of tight junction and gap junction proteins, ECM proteins, SLC influx transporters, ABC efflux transporters and metabolic enzymes to the barrier function of the BBB endothelium, a more detailed comparison and discussion of the expression of genes coding for these proteins in the three models is included below.

#### 2.3.1. Tight and gap junction proteins

The expression of VE-cadherin and CLDN5 in endothelial cell 2D monocultures and 5-cell organoids was initially demonstrated by immunocytochemistry (**Figures 1,3**). At this stage, we looked into the expression of genes of the *CLDN* family and *ESAM* that codes for the endothelial cell specific adhesion molecule (ESAM), a transmembrane junction protein with a similar structure to junctional adhesion molecules.^[29]^ 3-Cell organoids showed the overexpression of *CLDN5* which is by far the predominant CLDN in endothelium and that codes for the integral membrane tight junction protein CLDN5 and is a gatekeeper of neurological functions (**Figure 6A,F**).^[30]^ The expression of this gene was maximum in 3D endothelial cell monocultures. However, the number of endothelial cells in 3-cell organoids is smaller than that in endothelial cell monocultures. *CLDN1* and *CLDN12* were also expressed, though to a lower extent than in endothelial cell 2D monocultures (**Figure 6A, F**); *CLDN12* is not required for BBB tight junction function. In endothelial cell 2D monocultures, the expression of *CLDN* genes was in general lower than in both 3D systems except for *CLDN1, CLDN11* and *CLDN12* (**Figure 6 A, F**). In all the systems, the expression of *CLDN1, CLDN2, CLDN3, CLDN4, CLDN6, CLDN7, CLDN8, CLDN9, CLDN11, CLDN14, CLDN16, CLDN17, CLDN18* and *CLDN20* transcripts was relatively low (**Figure 6A,F**); these genes are more specific of epithelial (and not endothelial) tight junctions.^[30,31]^ A relatively high level of *CLDN15* could be observed in the endothelial cell monocultures, while *CLDN19* expression was detected in the 2D endothelial cell model, but not in 3D organoids (**Figure 6A,B**). *ESAM* was also expressed at a lower level in 3-cell organoids than in endothelial cell 2D monocultures (**Figure 6A,F**) because endothelial cells of mesoderm origin selectively encode the immunoglobulin family adhesion molecule ESAM, which mediates cell–cell adhesion through homophilic interactions.^[32]^ Other genes upregulated in 3-cell organoids with respect to endothelial cell 2D and 3D monocultures are *GJA1* that codes for the gap junction alpha-1 protein (GJA1) also known as connexin-43 (**Supplementary Table S3**);^[33]^ connexin hemichannels and gap junctions contribute to maintain the physiology of the BBB, participate in paracrine communication, and mediate efficient and rapid bidirectional inter-cellular transmission of electrical and chemical signals. Similarly, we found the upregulation of *VCAM1* that codes for the vascular cell adhesion molecule-1 (VCAM-1) protein, which mediates endothelial cell adhesion and *VWF* that codes for the von Willebrand factor (VWF), a glycoprotein that might be involved in brain homeostasis (**Supplementary Table S3**).^[34]^

#### 2.3.2. Extracellular matrix proteins

The ECM consists of multimeric proteins and proteoglycans that participate in cellular migration, differentiation, and function as a support system for endothelial cells and astrocytes. The ECM is pivotal for development, function, and regulation of vasculature, tight junctions, neurons, and astrocytes through cellular signalling and adhesion.^[35]^ Lack of any ECM component can result in developmental and functional flaws. Structurally, the basement membrane (BM) is a highly organised protein sheet with a thickness of 50–100 nm. Biochemically, the BM consists of four major ECM proteins: collagen type IV, laminin, nidogen and perlecan (**Supplementary Figure S6**). In this context, we analyzed genes coding for key BM proteins.

Collagen type IV is the most abundant component of the BM. The α-chain of this protein consists of three domains and it is thought that six α-chains self-assemble into triple-helical molecules and form spider web-like scaffolds that interact with laminin.^[36]^ *COL4A1* and *COL4A4* that code for collagen IV α-chains are upregulated in our 3-human cell organoids with respect to endothelial cell monocultures (**Figure 6B,G**); COL4A1 is a highly conserved protein across species and is involved in angiogenesis.

Laminin is a T-/cruciform-shaped trimeric protein composed of α, β and γ chains. Brain endothelial cells, pericytes and astrocytes produce different isoforms of laminin at the BBB. *LAMA1, LAMA2* and *LAMA4* (coding for laminin α1, α2 and α4, respectively), which regulate the maturation and function of the BBB, and *LAMB2* and *LAMC3* (coding for β and γ chains of laminin) were also upregulated in the multicellular organoids (**Figure 6B,G**). Other genes showing higher in 3-human cell organoids than in endothelial cell monocultures are *HSPG2* (coding for perlecan, the core protein of the glycosaminoglycan heparin sulfate) and *NID1* and *NID2* (coding for nidogen-1 and 2, respectively) that serve as linker for collagen IV, laminin and other ECM proteins (**Figure 6B,G**).

#### 2.3.3. Active efflux transporters

ABC transporters are highly expressed by the BBB endothelium and they play a key role in maintaining the brain homeostasis because they actively govern the entry of compounds from the bloodstream into the CNS.^[37]^ As expected, 3-cell organoids expressed a moderately higher amount of genes coding for P-glycoprotein (*ABCB1*), and several multidrug resistance proteins (MRPs) from the C subfamily (*ABCC3, ABCC4, ABCC5, ABCC10*, and *ABCC11*) (**Figure 6C,H**); it is noteworthy that in this system only ∼1/3 of the cellular component in the multicellular spheroids is endothelial cells. MRP3 is a glycoprotein with a similar molecular mass as MRP2, with similar amino acid composition, and with overlapping substrate specificity. Human MRP3 is the only basolateral efflux pump shown to transport bilirubin glucuronides. In some cases such as *MRP2* (*ABCC2*) deficiency, *MRP3* (*ABCC3*) is strongly upregulated (**Figure 6C,H**). MRPs 1, 2, 7, 8 genes were not substantially expressed in any of the models (**Figure 6C,H**).

The alanine, serine, and cysteine transporters belong to the SLC1A family of excitatory amino acid transporters (EAAT), Na^+^-dependent proteins that reside in the membrane of astrocytes, neurons and the abluminal (brain-facing) membrane of the BBB.^[38]^ EAATs are involved in the efflux transport of glutamate across the BBB and ensure low levels of this neurotransmitter in the interstitial fluid of the brain. Genes coding for EAAT1 (SLC1A3, GLAST), EAAT2 (SLC1A2, GLT1), EAAT3 (SLC1A1, EAAC1) and EAAT5 (SLC1A7) are upregulated in 3-cell organoids with respect to 2D and 3D endothelial cell monocultures (**Figure 6D,I**), most probably due to the contribution of hAs to the total expression. These results were in good agreement with previous works that showed their expression in endothelial cells isolated from brain capillaries.^[39]^

#### 2.3.4. Active influx transporters

Different influx transporters are involved in the transport of essential endogenous nutrients (e.g., amino acids, glucose) from the bloodstream into the CNS and are critical for the normal function of the brain.^[40]^ Endothelial cell 3D monocultures expressed relatively high levels of *SLC2A3* (*GLUT3*) encoding for glucose transporter-3 (GLUT3) (**Figure 6D,I**), a pump that is more characteristic of all neurons,^[41]^ and did not express *SLC2A5* (*GLUT5*) that codes for the transporter GLUT5 (characteristic of enterocytes).^[42]^ Our 3-cell organoids expressed high levels of genes coding for different glucose transporters (**Figure 6D,I**). Of special interest is *SLC2A1* (*GLUT1*) that codes GLUT1 which is crucial for the development of the cerebral microvasculature with BBB properties *in vivo*.^[43]^ Glucose is the predominant energy source for the brain and heart; therefore brain is the most energy-demanding organ in which endothelium and astrocytes plays a major role in regulating their metabolism.^[44]^ The transport of glucose across the BBB into the brain is mediated by the facilitative glucose transporter GLUT-1. This gene was upregulated in the 3-cell organoids with respect to endothelial cell 3D and 2D monocultures, which constitutes another confirmation of the more physiological phenotype of the endothelium in our multicellular model.

Other genes that were highly expressed in 3-cell organoids when compared to 2D and 3D endothelial cell monocultures are *SLC16A2* (*MCT2*) and *SLC16A6 (MCT6*) that code for proton-coupled monocarboxylic acid transporters (MCTs) and *SLC6A6* coding for Na^+^- and Cl^-^-dependent taurine transporter (TauT) (**Figure 6D,J**). TauT plays a key role in many biological pathways such as neurotransmission. *SLC6A1*, the gene coding for the voltage-dependent gamma-aminobutyric acid (GABA) transporter SLC6A1 was also upregulated in the 3-cell organoids. This transporter is responsible for the re-uptake of GABA from the synapse; GABA counterbalances neuronal excitation in the brain and any disruption of this balance may result in seizures. A similar trend was observed for *ABCA2* and *ABCA8*, coding for ABCA2, an endo-lysosomal protein that plays an important role in the homeostasis of various lipids and Alzheimer’s disease, and ABCA8 that regulates the lipid metabolism and is implicated in various CNS pathologies (**Figure 6E,J**).^[45]^ *SLC27A1* coding for the long-chain fatty acid transport protein 1 (FATP1) was also overexpressed in the 3-cell models (**Figure 6D,I**).

#### 2.3.5. Metabolic enzymes

Metabolizing enzymes in the BBB have a functional role in the local metabolism of drugs and other xenobiotics.^[46]^ Thus, the overexpression of genes coding for them is an additional proof of the more physiological behaviour of an *in vitro* cellular model. 3-Cell organoids expressed high levels of cytochrome P450 genes including *CYP2D6* and *CYP2R1* that were low in 2D and 3D endothelial cell monocultures (**Figure 6E,J**). Conversely, the expression of genes such as *GSTP1, SULT1A*, and *UGT1A1* coding for phase-II metabolic enzymes glutathione S-transferase π, sulfotransferase 1A1 and UDP-glucuronosyltransferase, respectively, was low in all the specimens (**Figure 6E,J**). In addition, *SOD1*, a gene coding for the apoptotic enzyme superoxide dismutase 1 (SOD1), and *GTPBP10* for a mitochondrial protein were downregulated in 3-cell organoids, whereas upregulated in endothelial cell 2D cultures (**Figure 6E,J**). These results indicate that our organoids do not exhibit hypoxic conditions. Overall, the comprehensive RNA-Seq analysis of our bio-fabricated 3-cell organoids confirmed high relative expression of endothelial cell specific genes involved in key signalling pathways that contribute to the establishment of a functional BBB.

### 2.4. Interaction of nanoparticles with 5-cell organoids

Our previous investigations conducted with primary rat cells suggested that neither forebrain nor olfactory neurons internalize polymeric nanoparticles.^[17]^ Conversely, they were internalized by primary microglia. These studies were conducted in 2D monocultures. Cell-cell connections between different components of the CNS, including neurons and microglia, in 3D may contribute to generate a more physiological milieu, to the increase the integrity and the function of the BBB endothelium and reciprocally affect the phenotype of the other cells actively implicated in the interaction with particulate matter.^[47]^

The transport of nanomaterials across the BBB endothelium is usually by transcytosis and is initiated by endocytosis, for which the size should be ≤200 nm.^[48]^ Depending on the shape and surface properties, particles as large at 500 nm could be transported to a more limited extent.

A main limitation of the existing *in vitro* models is that in general they do not include CNS macrophages. Upon characterization of our multicellular organoids by immunocytochemistry, electron microscopy and RNA-Seq, and the confirmation that the recapitulate a more physiological behavior, we studied the ability of different polymeric, ceramic and metallic nanoparticles to cross the outer endothelial cell monolayer and reach the organoid bulk. The properties of the NPs used in this work are summarized in **Supplementary Table S4**. Before use, NPs were diluted under sterile conditions and mixed with the corresponding culture medium to the final desired concentration.

#### 2.4.1. Polymeric nanoparticles

In this work, we used four polymeric NPs produced by the self-assembly of chitosan (CS)-,^[49]^ poly(vinyl alcohol) (PVA)-^[50]^ and hydrolyzed galactomannan (hGM)-based graft copolymers^[51]^ synthesized by the hydrophobization of the polymer backbone with poly(methyl methacrylate) (PMMA) and displaying size between 92 ± 4 to 463 ± 73 nm and from positive to negative Z-potential (**Supplementary Table S4**); these two properties govern the interaction of nanoparticulate matter with cells^[52]^ and were measured immediately before the biological experiments.

We hypothesize that owing to the cellular heterogeneity of the multicellular organoids, some immunocompetent cells (e.g., microglia) could be more susceptible to damage or, conversely, to uptake the NPs to a greater extent than others (e.g., neurons), as we previously demonstrated in 2D culture (31). Differential cell compatibility and uptake by BBB cells has been extensively documented in the literature. However, the effect of microglia in 3D multicellular systems has been never investigated before. To address these questions, polymeric NPs were fluorescently labeled with FITC or RITC and their interaction (e.g., permeability) with 5-cell organoids after 24 h of exposure characterized by CLSFM and LSFM. In general, studies revealed that 0.1% w/v NPs do not cause any morphological damage in the organoids.

When 5-cell organoids were exposed to crosslinked mixed CS-PMMA30:PVA-PMMA17 NPs, most of them accumulated on the organoid surface and a smaller fraction could be found inside it, as shown in **Figure 7A,B** by 2D and 2.5D CLSFM. However, cross-sectional CLSFM images cannot provide complete multi-view volumetric information of 3D organoids for which we need to detect the fluorescence intensity of each individual voxel. Cell uptake specificity was investigated by advanced LSFM. Images taken from different angles confirmed that, as opposed to CLSFM, some NPs permeate into the organoids and suggested the possible involvement of astroglia or microglia in the transport (**Figure 7C,D**). In case of mild injury/disturbance, astrocytes become phagocytes which remove “foreign” material and produce anti-inflammatory cytokines. Conversely, under excessive injury/insult, “reactive” astrocytes produce proinflammatory cytokines that recruit and activate microglia.^[53]^ Both pathways could be involved in the uptake of the nanoparticles into the organoids bulk. Similar results were observed with fluorescently labeled CS-PMMA33 (**Figure 7E-H**), crosslinked PVA-PMMA17 (**Figure 7I-L**), and hGM-PMMA28 NPs (**Figure 7M-P**). Furthermore, representation of the cells as dots (**Figure 7D,H,L,P**) confirmed that these nanoparticles are not harmful to the cells and that the cell density was not majorly affected by exposure to the different NPs.

**Figure 7.**
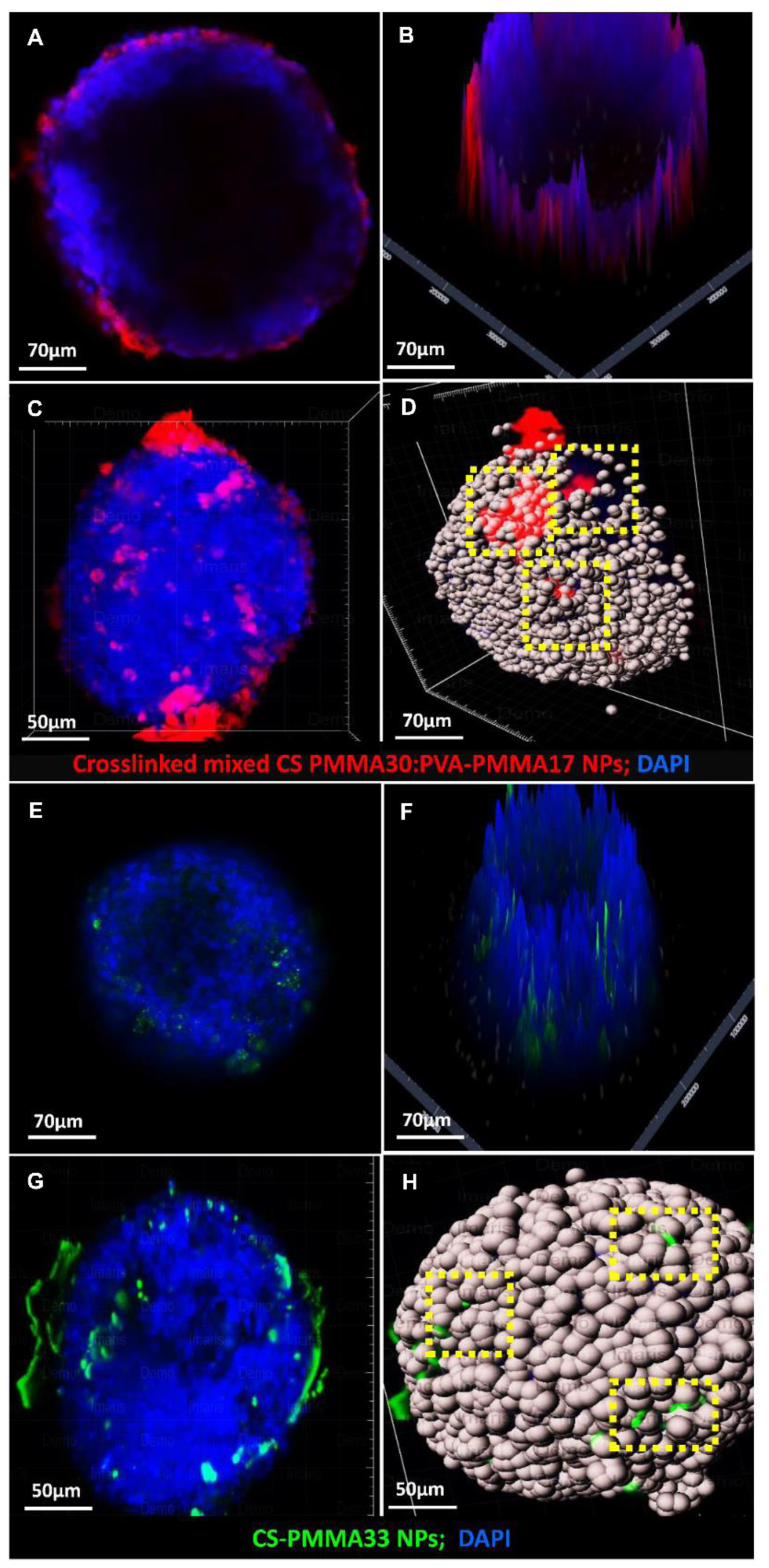
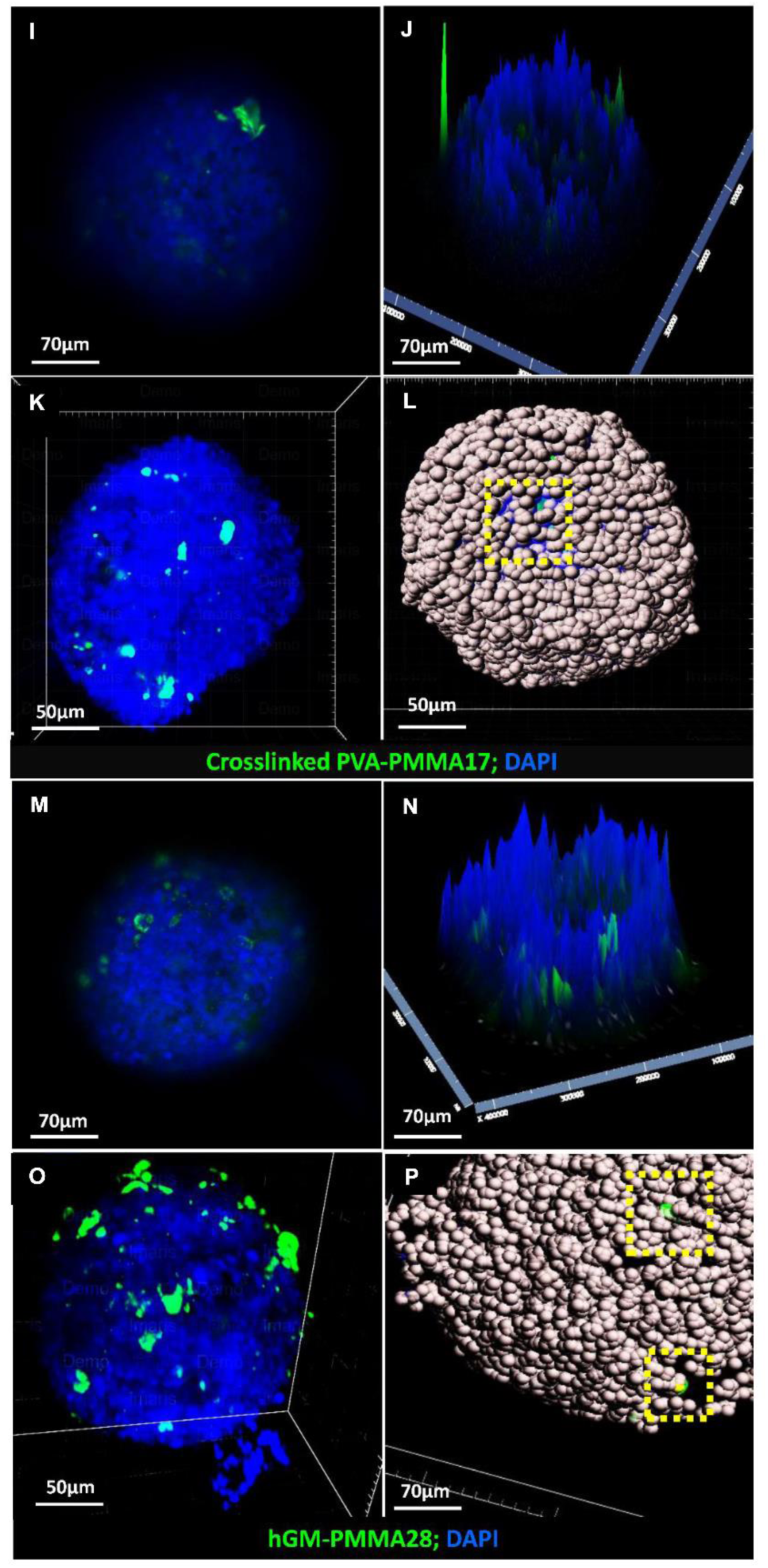
Characterization of the interaction of polymeric nanoparticles with bio-fabricated 5-cell neurovascular organoids. CLSFM micrographs of organoids exposed to (A,B) crosslinked CS-PMMA30:PVA-PMMA17, (E,F) crosslinked PVA-PMMA17, (I,J) CS-PMMA33, (M,N) hGM-PMMA28 nanoparticles. LSFM micrographs of organoids exposed to (C,D) crosslinked CS-PMMA30:PVA-PMMA17, (G,H) crosslinked PVA-PMMA17, (K,L) CS-PMMA33, (O,P) hGM-PMMA28 nanoparticles. Cell spots in D, H, L, and P are included to ease the visualization of the cells and exactly to locate the fluorescent nanoparticles underneath the organoid surface. Up-taken nanoparticles are highlighted by the dotted yellow squares.

Both fluorescence microscopic analyses confirmed that the enhanced fluorescence of NP-exposed organoids stemmed from NPs that most probably accumulated within endosomes/lysosome compartment of the primary microglia or immunocompetent astrocytes. Permeability studies conducted with our 5-cells organoids and metallic and carbon nanoparticles by STEM further contributed to our understanding of the possible uptake pathways (see below).

The brain is rich in energy-demanding nerve cells that metabolize glucose as the main fuel.^[54]^ Neurons consume glucose through glia cells, in which this nutrient is metabolized into lactate by the glycolytic pathway and transferred to axons and neuronal bodies when needed. To this end and to support their sentinel activity in the CNS, primary microglia overexpress GLUT1.^[55]^ In a previous work, we demonstrated that the accumulation of hGM-PMMA28 nanoparticles in pediatric sarcomas correlates well with the overexpression of GLUT1.^[51]^ Our LSFM results show that these inherently sugared nanoparticles are actively transported into the organoids, most probably by activated microglia and astrocytes. hGM-PMMA28 and other carbohydrate-based nanoparticles investigated in this work such as crosslinked mixed CS-PMMA30:PVA-PMMA17 and CS-PMMA33 could be also taken up through the mannose receptor that is expressed in microglia and astrocytes and that displays a carbohydrate recognition domain. Boric-acid crosslinked PVA-PMMA17 nanoparticles exhibit a boronated surface that may form complexes with microglia by toll-like receptor 4 (TLR4)–myeloid differentiation protein-2 (MD-2) signalling.^[56]^ In addition, microglia expresses sialic acid which may also bind boron.^[57]^ These findings are in good agreement with the active surveillance and homeostasis roles of microglia in the CNS microenvironment. Together with the gene expression pattern observed for endothelial cells cultured in 3-cell organoids with respect to 2D and 3D monocultures, this more complex multicellular *in vitro* model would recapitulate better the physiology of the BBB and serve as a platform to assess the interaction of nanomedicines and nano-pollutants with the CNS.

#### 2.4.2. Metallic nanoparticles

The use of metallic nanoparticles in nanomedicine is broad and varied.^[58]^ For example, Au and Ag NPs have been proposed in anti-cancer therapy^[59]^ and, upon injection, they could cross the BBB from the systemic circulation, and reach the CNS.^[60]^ Several studies used Au NPs as shuttles and demonstrated that the smaller the size, the higher the permeability; e.g., 10-20 nm Au NPs resulted in the highest cellular distribution in the brain of mouse.^[61]^ A main drawback of Au NPs is the limited ability of the CNS to clear them and the potential neurotoxicity associated with their accumulation.

To assess the performance of our new multicellular model, we exposed 5-cell organoids (5 days old) to ultra-small Au NPs (10 ± 2 nm, concentration of 1 × 10^6^ NPs/mL, **Supplementary Table S4**) for 24 h and analyzed their possible endocytosis by STEM. After 24 h, Au NPs were readily taken up and distributed in the cytosol, and inside endosomes and lysosomes of primary microglia that were identified by the presence of multiple lipid droplets in the cytosol (**Figure 8A,B**). The uptake mechanism is most probably energy-dependent, as shown elsewhere.^[62]^ We reported the ultrastructural morphology of microglia phenotype and high accumulation of lipid bodies is linked to a surveillant state in which microglia actively monitors the surrounding environment.

**Figure 8.**
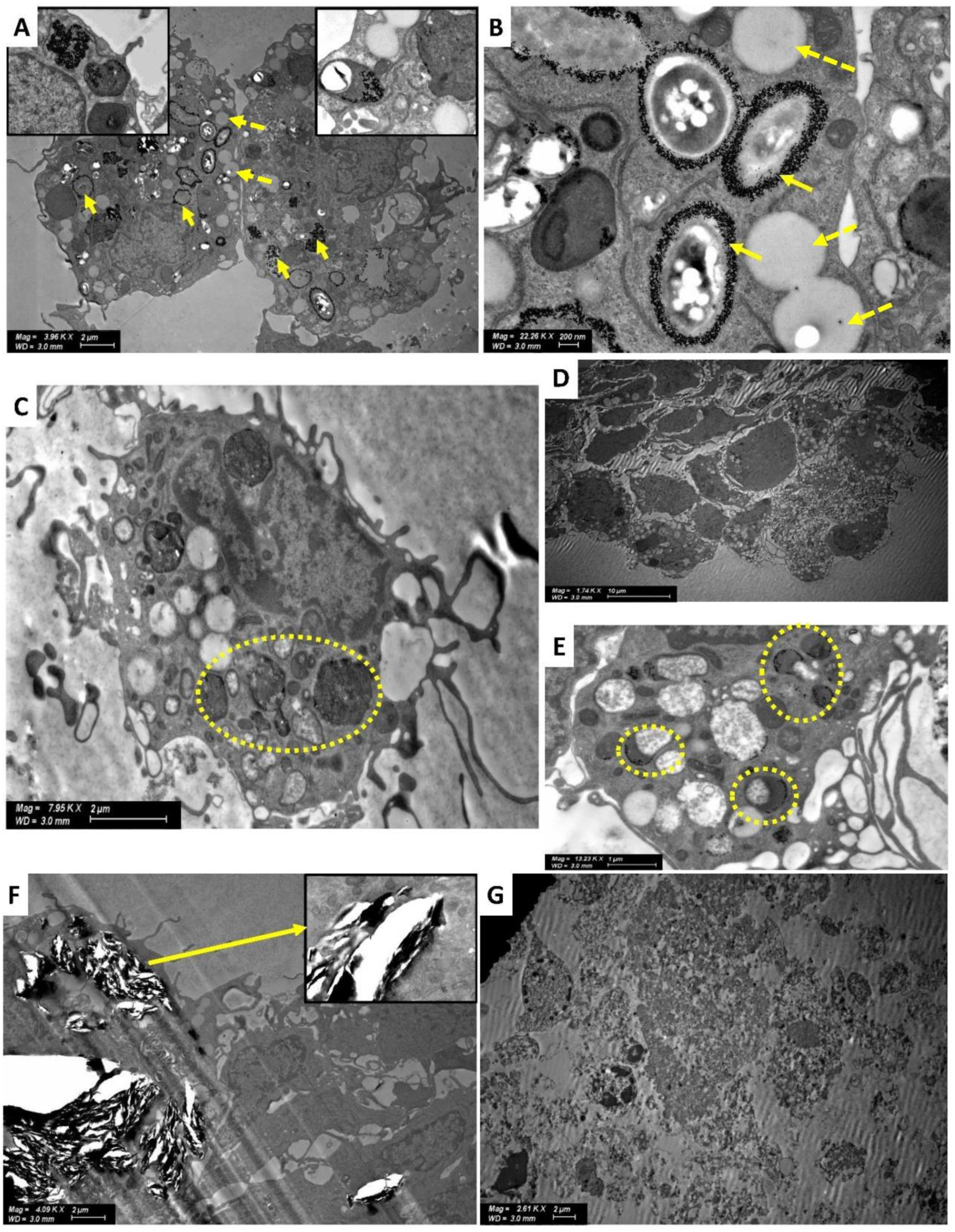
STEM micrographs of 5-cell organoids exposed to different metallic, carbon and ceramic nanoparticles. (A,B) Au NPs inside microglia (indicated with yellow arrows) and surrounding endosomes and lysosomes. Cells display characteristic lipid droplets (indicated with yellow dotted arrows). (C) Ag NPs inside vesicles (indicated with yellow dotted circle) of microglia displaying characteristic lipid droplets. (D) Ag NPs inside microglia clusters, (E) Ag NPs inside endosomes and lysosomes of glia (indicated with yellow dotted circle). (F) Cell disruption effect of graphene nanoplates on astrocytes (indicated with a yellow arrow in the insert). (G) Cell destructive effect of alkaline carbon dots on a 5-cells organoid.

Ag NPs have been also used in nanotherapeutics. Ag NPs (60 ± 13 nm, concentration of 1 × 10^6^ NPs/mL, **Supplementary Table S4**) were produced by a chemical method and organoids exposed to them for the same time. STEM studies focused on intracellular vesicles and revealed the presence of only very few Ag NPs inside these organoids (**Figure 8C-E**). This result might stem from the more limited cellular penetration of these NPs when compared to the much smaller Au counterparts. Another possible mechanism is that they undergo fast dissolution outside and inside the cells. These findings were consistent along different experiments.

#### 2.4.3. Graphene nanoplates and carbon dots

Among carbon nanomaterials, graphene-based ones are the most popular in the area of nanoneuroscience^[63]^ due to their various applications in neuronanotechnology^[63,64]^ and nanosafety studies.^[65]^ Here, we investigated the effect of graphene nanoplates (5 µm diameter and 10 nm thickness; 25 μg/mL, **Supplementary Table S4**) on our organoids. After 24 h, significant cell death could be observed, most probably due to direct cell damage (**Figure 8F**). Similarly, alkaline carbon dots (10 nm; 25 μg/mL, **Supplementary Table S4**) caused complete destruction of cellular structures (**Figure 8G**). In recent years, carbon dots have become a rising environmental concern because of their extensive use in energy applications, massive emissions to the air and likely transport from the nasal mucosa into the CNS. Our results strongly suggest the possible toxicity of these carbon NPs and the need for more systematic investigations that screen their detrimental effect on the CNS.

The interplay between other CNS-tissue-type cells, microglia and different nanomaterials by molecular level analysis which can provide sufficient sensibility to analyze the functional/phenogenotypic response of this brain’s own-macrophage cell population,^[66]^ was beyond the scope of the present work. In future research, this model will be characterized in other molecular level and individual cell specific aspects.

## 3. Conclusions

Several approaches have been utilized to develop preclinically and clinically relevant *in vitro* BBB models where human endothelial cells wrap up onto the surface of neural cell organoids. However, these models do not comprise a fundamental cellular player involved in the transport and elimination of particulate matter, microglia, and thus, their relevance in the study of the interaction of the CNS and nanomedicine and nano-pollutants is questioned. Previous models made of induced pluripotent stem cells showed that these cells cannot undergo differentiation into a complete set of neurovascular cells simultaneously and other cell types need to be added to create the complete brain environment.

To overcome the limitations in the field, in this work, we developed a multicellular organoid that combined human endothelial cells, pericytes and astrocytes with primary neurons and microglia isolated from rat brain in a ratio that mimics the cell composition of the brain tissue. The bio-fabricated organoids formed a layer of endothelial cells on the surface that formed adherens and tight junctions, two structures that govern paracellular permeability of molecules and particles at the BBB and thus, their bioavailability in the CNS. They also express active efflux transporters which are associated with the low bioavailability of neurotherapeutics in the brain. The organoids are stable in culture for at least two weeks and after comprehensive characterization, they were used to study the toxicity and permeability of different types of NPs. All the model polymeric NPs show good compatibility and undergo uptake. Similar results were observed with metallic nanoparticles, though the permeability depended on their size. In contrast, carbon NPs showed high toxicity, causing extensive cell death. The proposed multicellular organoid is easily bio-fabricated, and scalable to high-throughput capacity due to the simplicity of the method and the small number of reagents required. Furthermore, the screening throughput of this model can be increased even further through the possibility of integration with automated microscopy and robotics technologies. The ease of culture, cost-effectiveness and reproducibility of this model offer a very practical and attractive approach for researchers interested in studying BBB drug transport and developing brain-penetrant drugs for the treatment of CNS diseases. Taken together, our findings confirm that these multicellular systems form a functional BBB. Our model may serve as a valuable tool not only in nanotherapeutics, but in nanoneurosafety and nanoneurotoxicity.

## 4. Materials and Methods

### 4.1. Cell cultures

The hCMEC/D3 cell line derived from human temporal lobe microvessels (hBMECs, EMD Millipore, Burlington, MA, USA) was maintained in EndoGRO Basal Medium (EMD Millipore) with supplements containing 5% fetal bovine serum (FBS), L-glutamine, vitamin C, heparin sulfate and recombinant human epidermal growth factor (rhEGF), all purchased from Sigma-Aldrich (St. Louis, MO, USA).^[67]^ hAs (ScienCell Research Laboratories, Carlsbad, CA, USA) were grown in astrocyte growth medium (ScienCell Research Laboratories) containing 2% FBS supplemented with astrocyte growth factors, penicillin-streptomycin.^[68]^ hBVPs (ScienCell Research Laboratories) were maintained in pericyte culture medium (ScienCell Research Laboratories) containing 2% FBS, pericyte growth supplement and penicillin-streptomycin.^[69]^ All cells were incubated at 37°C in humidified 5% CO_2_/95% air.

### 4.2. Animals and isolation of primary neural-tissue cells

The use of neonate Sprague-Dawley (SD) rats (P0-1) and the protocols utilized for the isolation of the primary CNS cells were conducted with the approval of the Animal Care Committee of the Technion-Israel Institute of Technology. All the experimental procedures were in accordance with the guidelines set by the EU Council Directive (86/609 EEC) and according to the official protocol #IL-141-10-17 (expiry date 19 December 2021).

Primary neural, and neural stem/progenitor cells^[17, 70]^ were isolated from P0-1 SD rats as described below. Briefly, rats were sacrificed by decapitation, and whole brains were quickly removed aseptically (SZ09010122 Binocular Zoom Stereo Microscope, 8x∼50x, YSC Technologies, Fremont, CA, USA). Cerebral cortices of three pups were dissected under sterile conditions and kept on ice in a sterile Falcon tube (15 mL) containing 3 mL of trypsin-EDTA 0.25% (Sigma-Aldrich) for 10 min. The enzyme solution was aspirated, and enzyme-digested tissues were triturated in 10 mL of warm (37°C) Dulbecco’s Modified Eagle Medium (DMEM, Sigma-Aldrich) containing 10% heat-inactivated FBS. Cortical neuron cultures were dissociated by trituration using a 10 mL pipette (about 15-20 times) and cells were centrifuged (Z300 Hermle Micro Centrifuge, Hermle AG, Gosheim, Germany) at 4°C (1500g, 5 min) to obtain pellets. Single cells were separated from non-dissociated tissue debris by sieving them through Falcon™ Cell Strainers (pore diameter of 70μm, Sigma-Aldrich) and seeded onto 24- or 96-well plates coated with poly-L-lysine-(PLL, Sigma-Aldrich) at cell densities of 2 × 10^5^ or 1 × 10^4^ cells/well, respectively. Cultures were incubated at 37°C in a humidified atmosphere of 5% CO_2_/95% air. Cultured cells were grown in serum-free neuron-specific DMEM and Ham’s F12 nutrient mixture (DMEM: F12 1:1, Sigma-Aldrich) supplemented with 1% v/v insulin-transferrin-selenite (ITS, Sigma-Aldrich). For conventional cultures and bioimaging, the dissociated cultures were plated on glass coverslips in 24-well plates (Paul Marienfeld GmbH & Co. KG., Lauda-Königshofen, Germany). Before culture, glass coverslips were washed in water and 96% ethanol, air-dried, placed in 24-well plates, and coated with PLL overnight. The cell composition of the cultures was characterized by immunocytochemical staining for neuron specific biomarkers. Microglial cultures were prepared according to Saura *et al*.^[71]^ Briefly, P0-1 SD rat brains were removed and rinsed in phosphate buffer saline (PBS, Sigma-Aldrich). After careful removal of the meninges and brains cortices, they were mechanically dissociated and trypsinized for 20 min. Cells were cultured in 6-well plates with DMEM containing 10% FBS at 37°C in humidified 5% CO_2_/95% air. The medium was exchanged twice weekly. Microglia cells were isolated from mixed glia by mild trypsinization (30 min-2 h) with trypsin-EDTA 0.25% diluted 1:3 in serum-free DMEM, at 37°C. After detachment of astrocyte brown sheets, the firmly attached macrophages were further propagated in DMEM:F12 1:1 with 10% FBS and the cells replated in 24-well plates containing PLL-coated glass coverslips at a density of 50,000 cells/well (30). For microglia stainings, cells were fixed and the immunostaining and imaging were performed according to the protocol described below.

### 4.3. Assembly of multicellular organoids

Three-cell organoids comprising hCMEC/D3, hAs and hBVPs were cultured in EndoGRO Basal Medium supplemented with 5% FBS, L-glutamine, vitamin C, heparin sulfate and rhEGF, at 37°C in humidified 5% CO_2_/95% air using the liquid overlay culture system. We used the same method to produce 5-cell organoids incorporating primary neurons and microglia isolated from neonate rat (see above). This method is our adaptation of the aggregate cultures previously described.^[12a,12b,72]^ Briefly, the three or five cell types were harvested by trypsin-EDTA 0.25% and resuspended in ENDOGro culture medium. The concentration of hECs, hAs, hBVPs, primary neurons and microglia cells in each individual suspension was determined using a haemocytometer. Then, cells were resuspended in 100 μL of medium at a 4:2:1:1:1 ratio and transferred to ultralow attachment (ULA) round bottom 96-well plates (CellCarrier Spheroid ULA 96-well Microplates, PerkinElmer, Waltham, MA, USA). In another method, we prepared a 1% w/v agarose solution (molecular biology grade, Bio-Rad Laboratories, Hercules, CA, USA) in PBS under boiling until complete dissolution. The agarose solution (100 μL) was pipetted into each well of a 96-well plate, while it was still hot, and allowed to cool to 37°C and solidify. Then, a suspension of hCMEC/D3, hAs, hBVPs, and primary neurons and microglia cells at a 4:2:1:1:1 ratio was seeded in the 96-well plate. In both methods, cells were incubated in ENDOGro medium, at 37°C in humidified 5% CO_2_/95% air for 48–72 h to allow the formation of the spheroids.

To rule out possible detrimental human-rodent cell interactions during organoid assembly, hCMEC/D3 and primary rodent microglia cells were labeled with Cell Tracking Dye Kit - Green - Cytopainter (ab138891, Abcam, Cambridge, UK) and Cell Tracking Dye Kit - Deep Red - Cytopainter (ab138894, Abcam), respectively, and imaged by a GE IN Cell Analyzer 2000 imaging system (GE Healthcare, Chicago, IL, USA). For the staining, cells were mixed with each dye (1:1000 dilution in PBS) for 10 min, at 37°C in humidified 5% CO_2_/95% air, washed thrice with ENDOGro culture medium and co-cultured on CellCarrier Spheroid ULA 96-well Microplates to form 2-cell organoids, as described above. Images were acquired with a 10x objective at the end of each illumination period (every 10 min) for 12 h.

### 4.4. Characterization of multicellular organoids

#### 4.4.1. Immunofluorescent labeling

Organoids and monocultures were collected at day 5, pooled into a 0.2 mL Eppendorf tube (Corning Inc., Corning, NY, USA), washed thrice (5 min) with PBS and fixed in 4% w/v paraformaldehyde (PFA, Sigma-Aldrich) for 15 min at room temperature (RT). Fixed organoids were washed twice in PBS, permeabilized with 0.1% w/v Triton X-100 (Sigma-Aldrich) solution in PBS and blocked with 1% w/v bovine serum albumin (BSA, Sigma-Aldrich) for 1 h. Then, organoids were incubated in mixtures of primary antibodies overnight at RT under constant rotation, washed thrice with PBS, incubated overnight at 4°C in a mixture of secondary antibodies, all diluted in blocking solutions. Finally, the remainders of secondary antibodies were washed with PBS before staining cell nuclei with Hoechst 33342 (Sigma-Aldrich).

Immunolabeled organoids were visualized by using a LSM 710 confocal laser scanning fluorescence microscope (Carl Zeiss AG, Oberkochen, Germany) equipped with 10x, 20x, 40× water 1.4 NA and 63× oil (numerical aperture:1.3) Carl Zeiss objectives (as indicated in the corresponding figure legends). We used 405, 488, 561 and 647 nm lasers, and the scanning was done in line serial mode, pixel size was 50 × 50 nm. Image stacks were obtained with Zen Elements software. Whole-organoid images were acquired using a 10× magnification objective and images were processed by using the Zen 100 software.

The following commercial primary antibodies were used at a concentration of 1:500: (i) βIII-tubulin (AA10, sc-80016; Santa Cruz Biotechnology, Inc., Dallas TX, USA), (ii) glial fibrillary acidic protein (GFAP, 2E1, sc-33673, Santa Cruz), (iii) aquaporin-4 (AQP4; sc-390488 Santa Cruz), (iv) claudin-5 (CLDN5, sc-374221, Santa Cruz), (v) ionized calcium-binding adapter molecule 1/allograft inflammatory factor 1 (Iba-1/AIF-1, sc-32725, Santa Cruz), (vi) vascular endothelial (VE)-cadherin (ab33168, Abcam), (vii) inducible nitric acid synthase (iNOS, ab15323; Abcam), (viii) microtubule-associated proteins (MAP; sc-74421, Santa Cruz), and (ix) neuron-glial antigen-2 (NG2, sc-53389, Santa Cruz). The following commercial secondary antibodies were used at a concentration of 1:1000: (i) goat anti-rabbit IgG H&L (Alexa Fluor® 594, ab150080, Abcam), (ii) mouse IgG kappa binding protein (m-IgGκ BP) conjugated to CruzFluor™ 488 (sc-516176, Santa Cruz), (iii) mouse IgG kappa binding protein (m-IgGκ BP) conjugated to CruzFluor™ 555 (sc-516177, Santa Cruz), and (iv) mouse IgG kappa binding protein (m-IgGκ BP) conjugated to CruzFluor™ 647.

This method was also used to characterize the permeability of selected polymeric NPs and their localisation within the organoid. For this, NPs were synthesized by utilizing copolymers fluorescently-labeled by the conjugation of fluorescein isothiocyanate (FITC, green fluorescence, Sigma-Aldrich) or rhodamine isothiocyanate (RITC, red fluorescence Sigma-Aldrich) (30).

#### 4.4.2. Electron microscopy

Organoids were established for 24-72 h and collected and pooled in an Eppendorf tube. Then, they were washed once with PBS, and immersed in modified Karnovsky’s fixative (2% w/v glutaraldehyde and 3% w/v PFA in 0.1 M sodium cacodylate buffer containing 5mM CaCl_2_ and 3% sucrose, all from Sigma-Aldrich) for 1 h at RT. Organoids were washed in 0.1 M cacodylate buffer and fixed with 1% w/v osmium tetroxide (Sigma-Aldrich)/0.5% w/v potassium dichromate (1 h), stained with 1% w/v uranyl acetate (1h), washed twice with water and dehydrated in ethanol (Bio-Lab Ltd., Jerusalem, Israel) according to the following sequence: 50% (10 min), 70% (10 min), 90% (10 min) and 100% (2 × 10 min). Samples were embedded in EMbed 812 resin (Electron Microscopy Sciences, Hatfield, PA, USA) and polymerized at 60 °C for 48 h.^[73]^ Ultrathin sections of ∼70 nm were produced in an ultra-microtome (Leica Biosystems, Buffalo Grove, IL, USA), transferred to copper grids (Sigma-Aldrich) and visualized by using a Zeiss Ultra-Plus FEG-SEM (Carl Zeiss AG) equipped with STEM detector at accelerating voltage of 30 kV. This method was also used to characterize the permeability and the intracellular fate of model metallic and ceramic NPs in the organoids and enabled the elucidation of ultrastructural details and mechanisms of NP– organoid interactions.

Organoids were also imaged by cryo-SEM (Zeiss Ultra-Plus SEM) equipped with a Schottky field-emission gun and with a BalTec VCT100 cold-stage maintained below 145°C at low accelerated voltages of 1–1.2 kV to avoid radiation damage, and kept working distances of 3–5 mm. Everhart Thornley (‘‘SE2”) and the in-the column (“In Lens”) secondary electron imaging were used as detectors. For this, organoids were washed thrice with PBS and fixed by using high-pressure freezing (HPF). In the HPF technique, 1–3 µL of suspended organoids was sandwiched between two metal discs (3 mm diameter, 50 mm cavities), and high pressure (210 MPa) applied (High Pressure Machine HPM10, Bal-Tec AG, Balzers, Liechtenstein). This vitrification process prevents ice crystal formation and allows to prepare thicker samples with better preserved native ultrastructure.^[74]^ Frozen samples were mounted on a specialized sample table under liquid nitrogen and transferred by a high-vacuum cryo-shuttle (VCT100, Bal-Tec) to a Leica EM BAF060 freeze drying sample preparation system kept at 170°C for freeze-fracture with a cooled knife. Specimens were transferred under vacuum and at cryogenic temperature with the VCT100 precooled with liquid, for HR-Cryo-SEM imaging.

#### 4.4.3. Light sheet fluorescence microscopy

Organoids fixed in 4% w/v PFA and immunostained were embedded in 1% w/v low melting agarose (Bio-Rad Laboratories) solution and placed in an Eppendorf tube. Samples were analyzed by using a ZEISS Lightsheet Z.1 Fluorescence Microscope (Carl Zeiss) with the Sampler Starter Kit containing four color coded sleeves with their corresponding plungers to fit the glass capillaries high optical clarity for the 3D imaging to the sample holder. Upon selection of the proper capillary based on the sample size and plunger assembly, a 1% w/v low melting agarose solution (100 μL) in sterile PBS was mixed with the organoid, plunged into the capillary and allowed to polymerize at RT. Finally, the sample was pushed out and mounted in the sample chamber of the microscope at 37°C in 5% CO_2_/95% air for imaging. This method was also used to characterize the interaction of FITC-or RITC-labeled polymeric nanoparticles with the organoids. LSFM images were deconvolved using IMARIS Professional software (https://imaris.oxinst.com/).

#### 4.4.4. Total RNA sequencing and data processing

At the end of day 5, total RNA was extracted from BBB organoids using a T-series Ultraclear 1.5 ml Microcentrifuge tube (Scientific Specialties, Inc., Lodi, CA, USA). The organoid pellet was immersed in 500 µL of TRIzol (Invitrogen, Carlsbad, CA, USA) following the manufacturer’s instructions and stored at −80°C until further use. Samples were analysed in triplicates for RNA-Seq analysis. RNA-Seq libraries were constructed by using Illumina HiSeq 2500 Stranded Total RNA TruSeq RNA Library Preparation Kit v2 (Illumina, San Diego, CA, USA) and sequenced on the Illumina Nova Seq 6000 platform in the SR 50bp and barcode (Illumina). Total RNA-seq was performed from three replicates of 1-cell (only endothelial) and 5-cell organoids.

For gene expression analyses, trimmed reads with cutadapt were aligned to the reference genome (hg38UCSCassembly) using FASTQC version 0.11.5 (uses cutadapt version 1.10, Brabraham Bioinformatics, Cambridge, UK) and Tophat2 version 2.1.0 (uses Bowtie2 version 2.2.6, Johns Hopkins University, Baltimore, MD, USA), with default parameters and Ref Seq annotation (genome-build GRCh38.p9). The distribution of alignments was analysed using Cufflinks version 2.2.1 (Trapnell Lab, University of Washington, Seattle, WA, USA) and fragments per kilo base of exon model per million reads mapped values were quantile normalized. The quality control of the sequenced data (quality of all sequenced bases in all reads) was evaluated using FASTQC version 0.11.5. The quality scores are presented as Phred values (10 log10P base call is wrong), i.e. values higher than 30 indicate a probability of less than 10^−3^ of an incorrect base call (**Figures 1,2** of **Supplementary experimental file**). Only unique mapped reads were counted to genes, using ‘HTSeq-count’ package version 0.6.1 with ‘union’ mode. Normalization and differential expression analyses were conducted using DESeq2 R package version 1.10.0. The reads were mapped to the Human genome (ftp://ftp.ensembl.org/pub/release-96/fasta/homo_sapiens/) using Tophat2 version 2.1.0 (65), with up to 2 mismatches allowed per read, the minimum and maximum intron sizes were set to 70 and 50,000, respectively, and an annotation file was provided to the mapper (**Figure 3** of **Supplementary experimental file**).

#### 4.4.5. Protein expression analysis

5-Cell organoids were collected and washed with PBS several times and lysed by adding a NP-40 lysis buffer (Thermo Fisher Scientific) to conduct Western Blot analysis. Cell lysates were centrifuged (16,800g, 4°C) in a table-top centrifuge (Eppendorf, Hamburg, Germany) and stored at −80°C until use. Protein concentrations were determined by Bradford protein assay (Bio-Rad Laboratories) and SDS-PAGE was performed according to standard protocols. In each lane, total protein was loaded, and 10% or 15% SDS-PAGE gels were used. Gels were transferred to nitrocellulose membranes (GE Healthcare Life Sciences, Marlborough, MA, USA), blocked with 5% w/v non-fat milk (Sigma-Aldrich) for 1 h at RT, washed and incubated with the selective potentially associated primary antibody for overnight at 4°C (see above for antibodies list). Membranes were washed and incubated with the respective secondary HRP conjugate antibody (Bio-Rad Laboratories) for chemiluminescence analysis with an Image Quant LAS 4000 camera (GE Healthcare Life Sciences). The following primary antibodies were used at a concentration of 1:500: (i) βIII-tubulin antibody, (ii) GFAP, (iii) Iba-1/AIF-1 and (iv) VE-cadherin.

### 4.5. The nanoparticles

In this work, we utilized different polymeric, metallic and carbon nanoparticles to assess their permeability in 5-cell organoids.

#### 4.5.1. Polymeric nanoparticles

Different polymeric NPs were obtained by the self-assembly of amphiphilic graft copolymers of CS, PVA and hGM with PMMA. The copolymers were produced by the free radical graft polymerization of methyl methacrylate initiated by cerium(IV) ammonium nitrated in acid aqueous medium, as reported elsewhere.^[49b, 51]^

#### 4.5.2. Metallic nanoparticles

Au NPs (diameter of 10 nm, 741957; Sigma-Aldrich) and Ag NPs (diameter of 60 nm) synthesized from silver nitrate by reduction with sodium borohydride and stabilization with sodium citrate (all supplied by Sigma-Aldrich)^[75]^ were utilized as prototypes of metallic NPs.

#### 4.5.3. Carbon nanoparticles

We used graphene nanoplatelets (diameter of ∼5 µm and thickness of ∼100 nm, GNN P0205, Ants Ceramics Pvt. Ltd., Vasai-Virar, India) and alkaline carbon dots produced by the reaction of acetone with sodium hydroxide, as reported by Hou *et al*.^[76]^

The size (hydrodynamic diameter, D_*h*_) and size distribution (polydispersity, PDI) of the different nanoparticles before the biological studies were measured by dynamic light scattering (DLS, Zetasizer Nano-ZS, Malvern Instruments, Malvern, UK) in 10 mm quartz cuvettes using a He-Ne laser (673 nm) as light source at a scattering angle of 173°, at 25°C. Data were analyzed using CONTIN algorithms (Malvern Instruments). The surface charge was estimated by measuring the zeta-potential (Z-potential) by laser Doppler micro-electrophoresis in the Zetasizer Nano-ZS. Each value is expressed as mean ± S.D. of at least three independent samples, while each DLS or Z-potential measurement is an average of at least seven runs. For biological studies, copolymers were fluorescently-labeled with FITC or RITC, both from Sigma-Aldrich ^*[49b, 51]*^. The properties of the NPs used in this work are summarized in **Supplementary Table S1**. Before use, NPs were diluted under sterile conditions and mixed with the corresponding culture medium to the final desired concentration.

### 4.6. Statistical Analysis

RNA-Seq data to compare between organoid groups was obtained from three biological replicates. Wald test parameters were used for pairwise comparisons. The statistical analysis was conducted and the log2 fold change and the adjusted p-values (p_adj_) were indicated for significantly upregulated and downregulated genes; p_adj_ <0.05 was considered statistically significant. The differential expression analysis was conducted using ‘DESeq2’ R software. The Zen software was used to evaluate CLSFM and LSFM images. In some cases, the TIFF images acquired with IMARIS software were imported for subsequent measurement and analysis. Results are expressed as mean ± S.D. of three experimental replicates performed under the same conditions (n = 3), unless otherwise specified. All cellular measurements and subsequent analyses were performed in a blinded manner and cell counts and image processing for all experimental conditions were carried out in triplicate. CT values and normalized gene expression values were taken after analysis and the data were exported to Excel (Microsoft, Seattle, WA, USA) spreadsheet software for expression units and S.D. analysis.

## Supporting information

Supplementary experimental file

