## Supplementary Figures and Tables for "Multicellular Organoids of the Neurovascular Blood-Brain Barrier: A New Platform for Precision Neuronanomedicine"

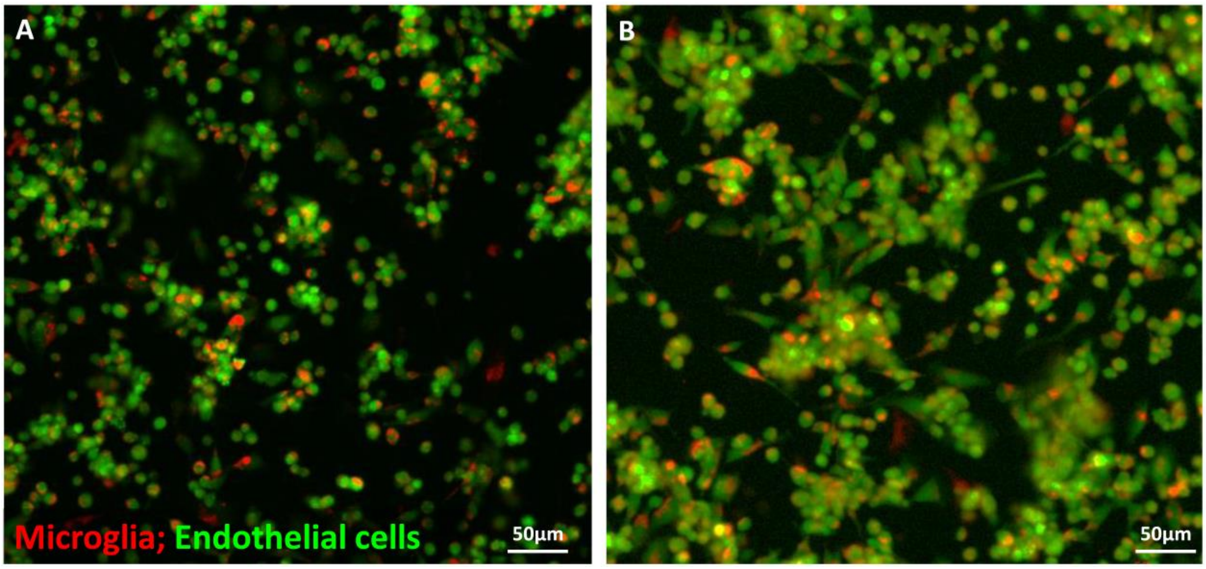


**Figure S1. Interspecies interaction of human and rat cells.** CLSFM micrographs of hCMEC/D3 endothelial cells (green) and primary rat microglia (red) stained with Cell Tracking Dye Kit - Green - Cytopainter and Cell Tracking Dye Kit - Deep Red – Cytopainter, respectively, after (A) 1 h and (B) 12 h.

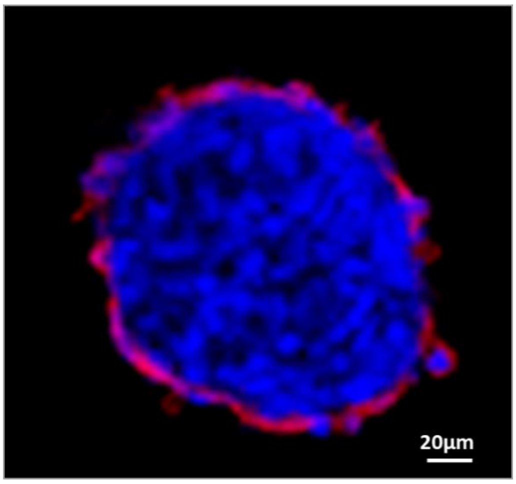


**Figure S2. Characterization of hCMEC/D3 endothelial cell distribution in 5-cell organoids.** CLSFM micrograph of an organoid showing endothelial cells immunostained for VE-cadherin (red) and nuclei of all the cells stained with DAPI (blue).

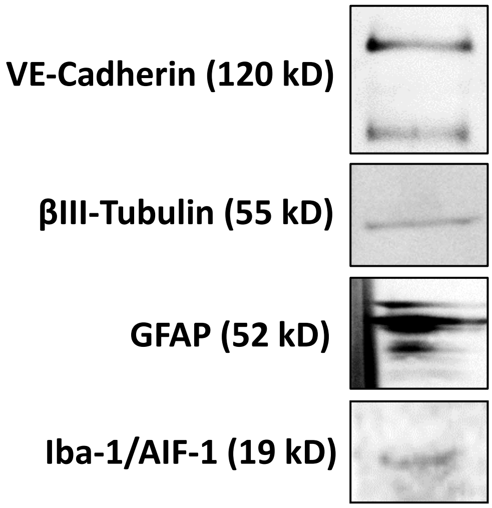


**Figure S3. Western blotting analysis of brain specific neurovascular organoids.** Confirmation of the expression of characteristic protein markers.

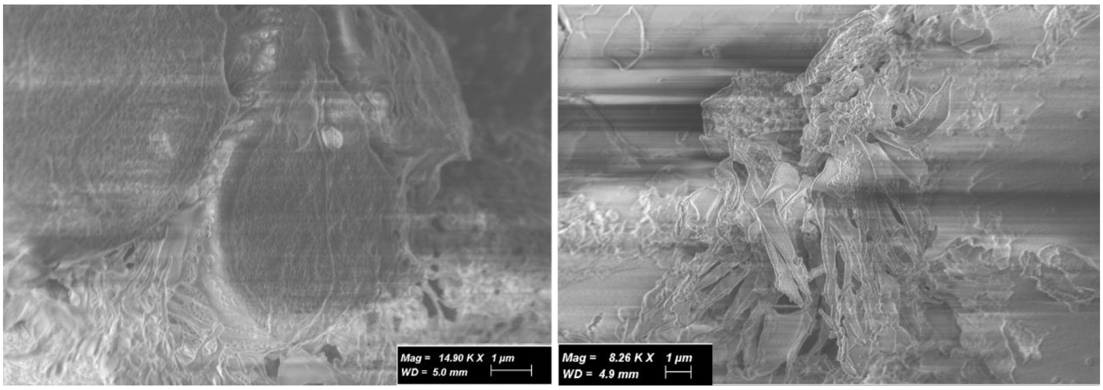


**Figure S4. Cryo-SEM tomogram of 3-cell organoids.**

**Table S1. Quality control of a high-quality total RNA sample by using 1% agarose gel electrophoresis.**

| Sample number | Sample name | Concentration, (ng/uL) [a] (Qubit®) | RIN (TS) |
|---|---|---|---|
| 1 | Endothelial cells 2D | 264 | 8.5 |
| 2 | Endothelial cells 2D | 116 | 10 |
| 3 | Endothelial cells 2D | 266 | 8.5 |
| 4 | Endothelial cells 3D | 110 | 9.6 |
| 5 | Endothelial cells 3D | 106 | 9.7 |
| 6 | Endothelial cells 3D | 111 | 9.6 |
| 7 | Human 3-cell organoid | 69.4 | 10 |
| 8 | Human 3-cell organoid | 99.4 | 10 |
| 9 | Human 3-cell organoid | 62.6 | 10 |

[a] Determined by Qubit®.

Note: RNA with excellent quality, RIN = 10; RNA with acceptable quality, RIN = 6.9; RNA with poor quality, RIN = 1.8.

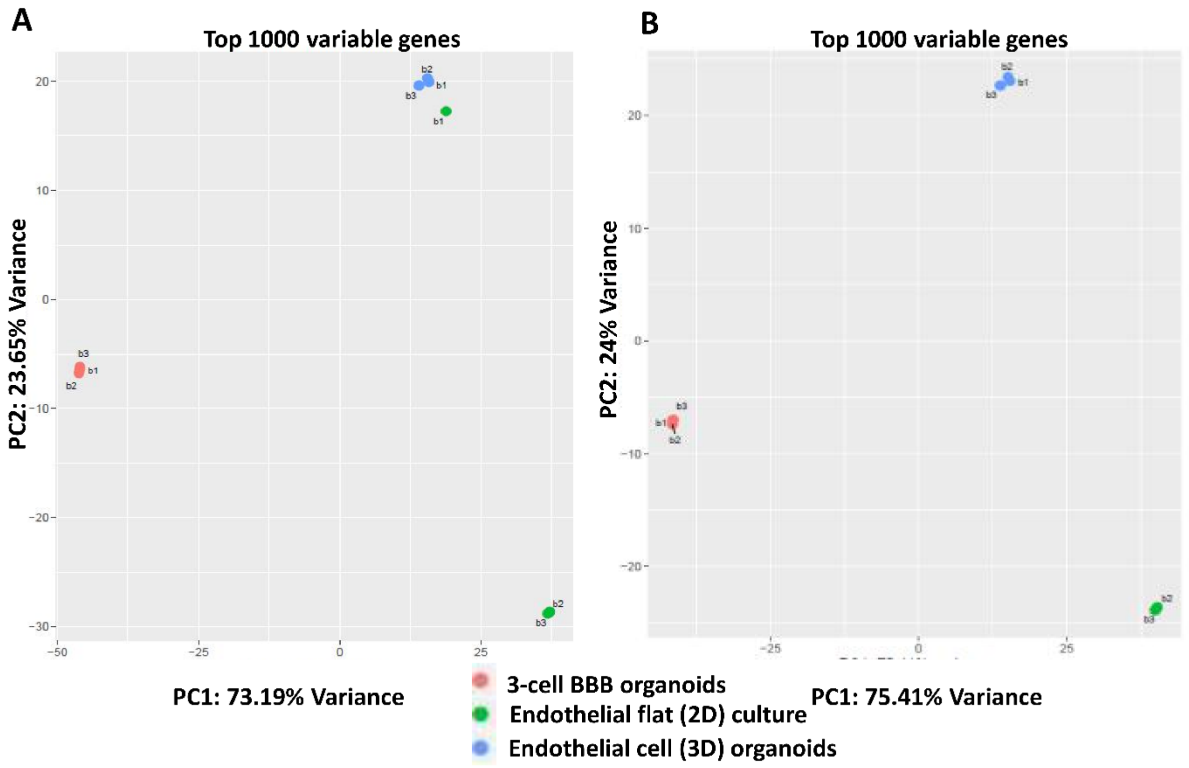


**Figure S5. Identification of genes that are most informative for defining cell subpopulations by PCA.** (A) PCA plot between the three groups and (B) PCA plot between groups, after excluding the outlier sample of endothelial flat (2D) culture.

**Table S2. Differential expression analysis of endothelial cells in 3-human cell organoids, endothelial cell (3D) organoids and endothelial cell flat (2D) cultures.** Pairwise testing between conditions.

| Comparison | Total # Genes | All Zero | Low Counts | Tested | Significant up-regulation | Significant down-regulation |
|---|---|---|---|---|---|---|
| 3-Human cell organoids vs._ endothelial cell flat (2D) cultures | 58,174 | 22,654 | 15,211 | 20,309 | 7314 | 6273 |
| 3-Human cell organoids vs._ endothelial cell (3D) organoids | 58,174 | 22,654 | 16,294 | 19,226 | 3966 | 3487 |
| Endothelial cell flat (2D) cultures vs._ endothelial cell (3D) organoids | 58,174 | 22,654 | 15,849 | 19,671 | 6290 | 6503 |

**Table S3. Comparative expression of different characteristic genes of endothelial cells in 3-human cell organoids, endothelial cell (3D) organoids and endothelial cell flat (2D) cultures, as determined by RNA-seq.**

| Gene | 3-Human cell organoids vs. endothelial cell flat (2D) cultures | 3-Human cell organoids vs. endothelial cell (3D) organoids | Endothelial cell flat (2D) cultures vs. endothelial cell (3D) organoids |
|---|---|---|---|
| *GJA1* | Upregulated | Upregulated | Downregulated |
| *EDN1* | Upregulated | Upregulated | Downregulated |
| *VWF* | Upregulated | Downregulated | Downregulated |
| *CD34* | Upregulated | Downregulated | Downregulated |
| *ENG* | Upregulated | Upregulated | Downregulated |
| *VCAM1* | Upregulated | No change | Downregulated |
| *EMCN* | Upregulated | Upregulated | No change |
| *NR2F2* | Upregulated | Upregulated | Downregulated |
| *CDH5* | Upregulated | Downregulated | Downregulated |
| *GJA4* | No change | No change | No change |
| *EPHB4* | Upregulated | Downregulated | Downregulated |
| *MCAM* | Upregulated | Upregulated | Downregulated |
| *FLT1* | No change | Downregulated | No change |
| *NOS3* | No change | No change | No change |
| *FLT4* | No change | No change | No change |
| *PTPRC* | No change | No change | No change |
| *CD4* | Upregulated | Upregulated | No change |
| *ICAM3* | No change | No change | No change |
| *BCL6* | Upregulated | Upregulated | Upregulated |
| *CD28* | No change | No change | No change |
| *ITGA4* | Up | Upregulated | Downregulated |
| *CD38* | No | Upregulated | No change |
| *CD86* | No | No change | No change |
| *MS4A1* | No | No change | No change |

**Table S4. Properties of the different polymeric, metallic and carbon nanoparticles used to investigate the interaction of nanomaterials with 5-cell organoids.**

| **Nanoparticle type** | **$D_h$ (nm) ± S.D.** | **PDI** | **Z-potential (mV)** | **Nanoparticle description** |
|---|---|---|---|---|
| **Polymeric** | | | | |
| CS-PMMA33 | 188 ± 9 | 0.30 | +23.0 | Non-crosslinked amphiphilic nanoparticles produced by the self-assembly of a graft copolymer of chitosan (CS) and poly(methyl methacrylate) (PMMA) containing 33% w/v of PMMA[49a] |
| Crosslinked mixed CS-PMMA30:PVA-PMMA17 | 463 ± 73 | 0.55 | +3.0 | Mixed amphiphilic nanoparticles produced by the self-assembly of a 1:1 weight ratio mixture of a graft copolymer of chitosan (CS) and poly(methyl methacrylate) (PMMA) containing 30% w/v of PMMA and a graft copolymer of poly(vinyl alcohol) (PVA) and PMMA containing 17% w/v of PMMA. Nanoparticles were ionotropically crosslinked with sodium |

| | | | | |
|---|---|---|---|---|
| | | | | tripolyphosphate which reacts with CS domains[49b] |
| Crosslinked PVA-PMMA17 | 92 ± 4 | 0.14 | -14.6 | Amphiphilic nanoparticles produced by the self-assembly of a graft copolymer of poly(vinyl alcohol) (PVA) and poly(methyl methacrylate) (PMMA) containing 17% w/v of PMMA (*1*). Nanoparticles were non-covalently crosslinked with boric acid which reacts with PVA domains[50] |
| hGM-PMMA28 | 141 ± 3 | 0.11 | -0.4 | Amphiphilic nanoparticles produced by the self-assembly of a graft copolymer of hydrolyzed galactomannan (hGM) and poly(methyl methacrylate) (PMMA) containing 28% w/v of PMMA[51] |
| **Metallic** | | | | |
| Silver | 60 ± 13 | | -38.0 | An aqueous solution of silver nitrate was heated to boiling, sodium citrate and sodium borohydride was added slowly[75a] |
| Gold | 10 ± 2 | <0.20 | -17.8 | - |
| **Ceramic** | | | | |
| Graphene nanoplatelets | ~5 µm diameter and 10 nm thickness | - | - | - |

| | | | | |
|---|---|---|---|---|
| Alkaline carbon dots | 10 ± 3 | 0.20 | −12.0 mV | *Aldol Reaction:* Sodium hydroxide was mixed with mL acetone under vigorous magnetic stirring for 1 h, and then the mixture was placed at ambient air, temperature, and pressure. The product was separated by centrifugation and washed to get CQDs powder[76] |

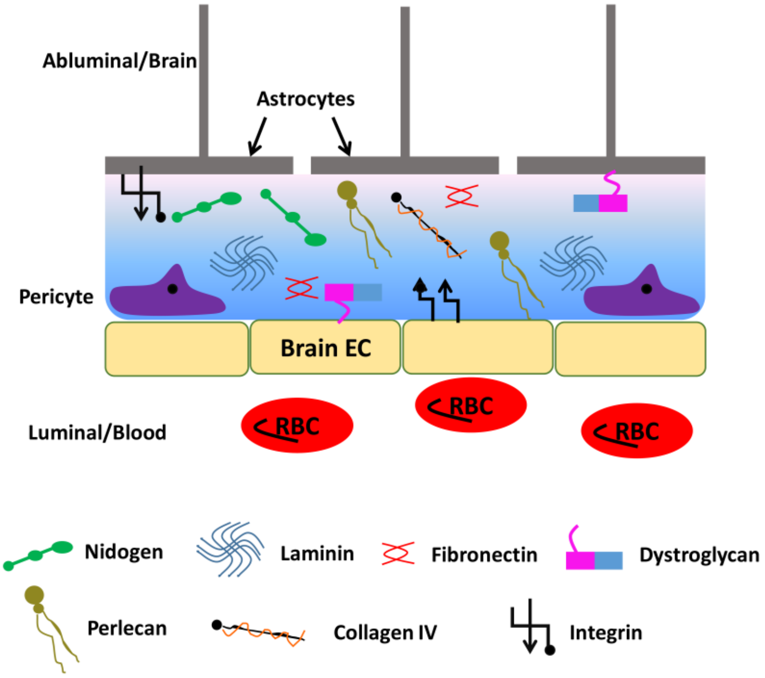


**Figure S6. Scheme of the extracellular matrix (ECM) in the vascular and parenchymal basement membrane of endothelial cells (EC) and red blood cells (RBC).** Key proteins are indicated.
