## Supplementary experimental file for "Multicellular Organoids of the Neurovascular Blood-Brain Barrier: A New Platform for Precision Neuronanomedicine"

### Analysis Pipeline

**Software and applications used:**

Library quality control: FASTQC version 0.11.5
Quality and adapter trimming: trim_galore (uses cutadapt version 1.10)
Mapping: Tophat2 version 2.1.0 (uses Bowtie2 version 2.2.6)
Gene counting: HTseq-count version 0.6.1/0.11.2
Normalization and differential expression analysis: DESeq2 R package version 1.18.1

**Input files**

The reference genome and the annotation file) used in the analysis was of *Human sapience GRCh38*, taken from ensembl repository:
ftp://ftp.ensembl.org/pub/release-96/fasta/homo_sapiens/
The reference GTF file was taken also form ensembl repository:
ftp://ftp.ensembl.org/pub/release-96/gtf/homo_sapiens/
All these files can be found in our FTP server, see section 4- Analysis result files.

**Table 1. Technical information and quality of sequencing.**

| Lane | Sample | Index | Quality of Run<br>Control - Mapped (%) | Control - Mismatch (%) | PF reads (%) | Unknown reads (%) | Number PF reads |
|---|---|---|---|---|---|---|---|
| 8 | BBB organoids | AGTCAA | | | | | 25,990,463 |
| | | AGTTCC | | | | | 24,495,706 |
| | | GTGGCC | | | | | 21,842,753 |
| | ECs 2D flat culture | ATGTCA | | | | | 25,571,133 |
| | | CCGTCC | 1.46 | 0.06 | 91.95 | 2.18 | 27,419,486 |
| | | GTTTCG | | | | | 26,094,770 |
| | EC 3D organoids | GTCCGC | | | | | 25,944,636 |
| | | GTGAAA | | | | | 25,979,188 |
| | | CGTACG | | | | | 22,945,300 |

- **Insert** – the DNA/cDNA sequence located between sequencing adapters.
  **Read** – one end of the paired-end segments.

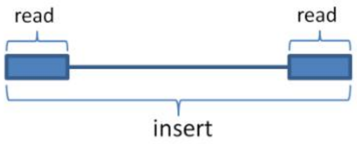


- **Control -** PhiX is a bacteriophage with a known genome sequence that is used as a standard sequencing control to estimate read accuracy. 1-2% of the reads of each lane are comprised of a PhiX sample which is mapped to the PhiX genome in order to estimate the error rate and quality of the sequencing run.

  A good control would have approximately 1% mapped control reads, and up to 1% mismatches (error rate).

- **Unknown reads -** an adapter containing a sample specific barcode is added to each library during library preparation. In some cases, if a read's sequenced barcode contains sequencing errors, then the read cannot be specifically identified with any sample and it is considered as “unknown”. The percentage reported in the table is out of the total number of reads per lane.
- **PF** (Passed Filter) **-** indicates reads that passed the automatic quality filter of the sequencer.

**Table 2. Trimming statistics.**

| Sample | Total reads | Containing adapter (%) Reads | Trimmed bases (%) | Reads removed (%) | Reads removed (#) |
|---|---|---|---|---|---|
| 3-cell BBB organoids | 25,990,463 | 2.6 | 0.3 | 0.1 | 18,349 |
| | 24,495,706 | 2.6 | 0.3 | 0.1 | 17,334 |
| | 21,842,753 | 2.6 | 0.3 | 0.1 | 17,212 |
| ECs 2D-culture | 25,571,133 | 2.6 | 0.3 | 0.1 | 17,660 |
| | 27,419,486 | 2.7 | 0.3 | 0.1 | 19,354 |
| | 26,094,770 | 2.7 | 0.3 | 0.1 | 17,324 |
| EC 3D-organoids | 25,944,636 | 2.7 | 0.3 | 0.1 | 24,616 |
| | 25,979,188 | 2.6 | 0.3 | 0.1 | 21,511 |
| | 22,945,300 | 2.6 | 0.3 | 0.1 | 16,927 |

The quality of the sequenced data (all sequenced bases in all reads) before and after trimming, as well as read length distributions after trimming, were evaluated using FASTQC. The quality scores are presented as Phred values (10log10P (*base call is wrong*)), i.e. values higher than 30 indicate a probability of less than $10^{-3}$ of an incorrect base call.

**Table 3. Mapping statistics.**

| **Sample** | **Total no reads** | % unmapped | % Uniquely unmapped | # Uniquely unmapped | % Multi Mapped | % Gapped Uniquely mapped reads (out of unique) |
|---|---|---|---|---|---|---|
| BBB organoids | 25,972,114 | 1.99 | 95.12 | 24,703,511 | 2.9 | 20.32 |
| | 24,478,372 | 2.05 | 95.12 | 23,283,509 | 2.83 | 19.99 |
| | 21,825,541 | 2 | 95.2 | 20,777,273 | 2.8 | 20.2 |
| ECs 2D flat culture | 25,553,473 | 1.89 | 95.28 | 24,347,151 | 2.84 | 20.65 |
| | 27,400,132 | 2.56 | 94.09 | 25,780,893 | 3.35 | 16.81 |
| | 26,077,446 | 2.64 | 93.87 | 24,479,978 | 3.48 | 16.64 |
| EC 3D organoids | 25,920,020 | 2.15 | 94.1 | 24,390,672 | 3.75 | 18.86 |
| | 25,957,677 | 2.09 | 94.65 | 24,570,220 | 3.26 | 19.04 |
| | 22,928,373 | 2.12 | 94.5 | 21,668,202 | 3.38 | 18.68 |

•**Uniquely mapped** – Reads aligned with high confidence to a single genomic location with up to 2 mismatches. Only the uniquely mapped reads are used for further analysis.
•**Unmapped** – Reads for which no alignment was found to the reference genome.
•**Multi-mapped** – Reads mapped to more than one possible location in the genome. These reads are not used in the analysis.
•**Gapped uniquely mapping** – Reads aligned uniquely to a splice junction **(Figure. 3)**

### Quality control and trimming:

The quality of the sequenced data (quality of all sequenced bases in all reads) was evaluated using FASTQC. The quality scores are presented as phred values (10log10P (*base call is wrong*)), i.e. values higher than 30 indicate a probability of less than $10^{-3}$ an incorrect base call.

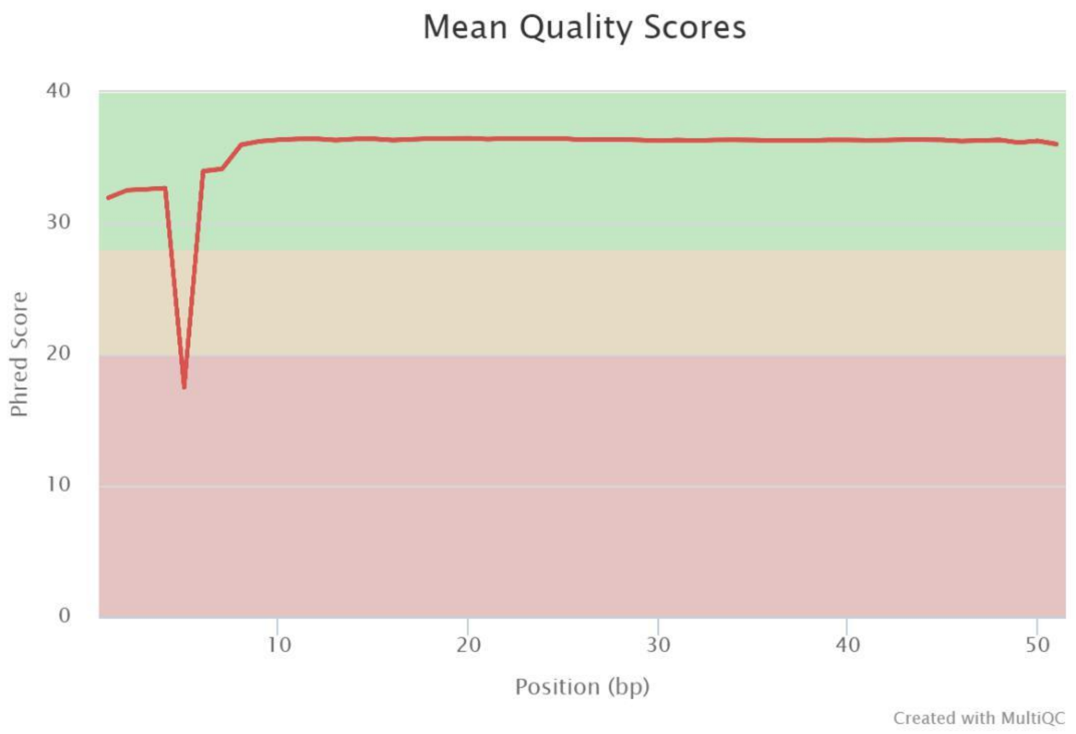


**Figure 1. Per-base quality scores.**

### Conclusions from the statistical data

The number and percentage of reads that passed the automatic sequencer filter in conjunction with the control mapping value, error rate, and per-base quality scores demonstrate that the run was of high quality.

### Quality control and trimming:

Adapter and quality trimming were performed on the data in order to optimize mapping accuracy.

The minimal quality threshold was set to 20 (phred scale) and the selected TruSeq RNA adapter sequence was: AGATCGGAAGAGC. The minimal length required for each read after trimming was set to 30 and only inserts for which both reads passed the filtering criteria were used.

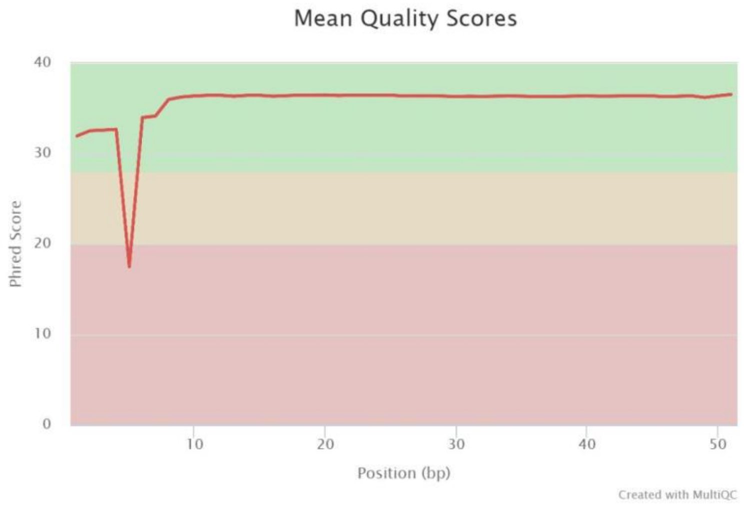


**Figure 2. Per-base quality.**

**Conclusions from the statistical data**

The amount and percentage of reads that passed the sequencer's automated quality filter, control mapping value, error rate, and per-base quality scores indicate that the sequencing was of high quality. Only a small percentage of reads were discarded due to trimming and the peak of the length distribution is at the original length of 51bp.

### A. Mapping

**Alignment details (Tophat):**

Tophat is a program designed specifically for alignment of RNA-Seq data to a chosen reference genome. The software uses the high-throughput, short read aligner 'Bowtie2' and analyzes the mapping results to identify splice junctions between exons.

Tophat divides the reads into shorter segments and generates a database of possible splice junctions (built using both the input data of the current project as well as a given annotation file specifying the coordinates of all known exons of the organism). Input reads are then mapped to the uniquely generated database of possible splice junctions.

TopHat parameters were specified for identification of alignments with a maximum of 3 mismatches from the reference sequence per read, and a maximum of 3 mismatches for a segment of a read. As per literature precedents, the minimum and maximum intron sizes were set to 70 and 500,000, respectively.

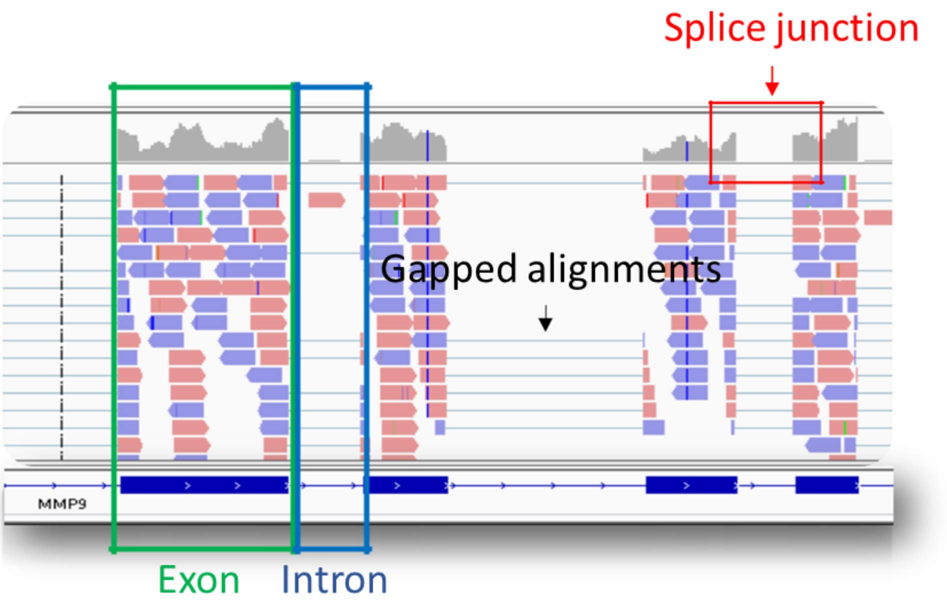


**Figure 3. Illustration of RNA-Seq read alignments using 'Tophat' – A graphic representation of the mapping patterns of RNA-Seq reads to a reference genome.** The figure above reveals the distribution of reads that are mapped to exons (outlined in green), a splice junction (outlined in red), and introns (outlined in blue). The figure allows the user to see that the majority of reads map to exons, slightly fewer reads map in a gapped manner to the pictured splice junction, and—consistent with the nature of RNA library preparation preference for mature mRNAs—almost no reads map to introns.

### B. Raw gene counts:

Uniquely mapped reads are assigned to genes based on the gene annotation file (specifying the coordinates of all exons of the annotated genes). The 'HTSeq-count' package was used to obtain gene counts with the "union" mode option and the parameter '--stranded' was set to 'no' according to the sample preparation kit specifications. For more information, please see the HTSeq-count documentation: http://www-huber.embl.de/users/anders/HTSeq/doc/count.html. The table below shows the distribution of reads.

**Table 4. Read assignments according to gene annotations.**

| Sample | **# reads unique** | % reads unique | # no feature | % no feature | Number ambiguous | %ambiguous |
|---|---|---|---|---|---|---|
| BBB organoids | 21,478,458 | 86.94 | 1,734,480 | 7.02 | 1,490,573 | 6.03 |
| | 20,218,632 | 86.84 | 1,679,234 | 7.21 | 1,385,643 | 5.95 |
| | 18,066,504 | 86.95 | 1,453,586 | 7 | 1,257,183 | 6.05 |
| ECs 2D flat culture | 21,637,878 | 88.87 | 1,182,240 | 4.86 | 1,527,033 | 6.27 |
| | 21,594,711 | 83.76 | 2,686,051 | 10.42 | 1,500,131 | 5.82 |
| | 20,471,953 | 83.63 | 2,580,918 | 10.54 | 1,427,107 | 5.83 |
| EC 3D organoids | 20,884,278 | 85.62 | 2,006,406 | 8.23 | 1,499,988 | 6.15 |
| | 21,168,497 | 86.16 | 1,863,925 | 7.59 | 1,537,798 | 6.26 |
| | 18,520,642 | 85.47 | 1,809,475 | 8.35 | 1,338,085 | 6.18 |

•**Counted reads** – uniquely mapped reads assigned to annotated exons.
•**No feature** – reads that could not be assigned to any annotated gene.
•**Ambiguous –** reads that can be assigned to more than one annotated gene and are therefore not counted to any gene.

### C. Gene counts normalization:

Normalization of raw counts is an important step which brings all samples to a common scale and enables comparing expression levels between different biological conditions. The need for normalizations may arise due to several possible reasons. First being technical deviations, such as different library sizes (total number of reads) when sequencing several samples together on a lane. Another possible cause is the biological condition, for example a certain treatment may cause a significant up-regulation of a small subset of genes, and thus affecting the overall reads distribution among genes. Normalization is conducted using 'DESeq2' R package (which is also used for the differential expression analysis), during the process a normalization factor (size factor) is calculated for each sample and the raw counts are divided by this factor. For details look at the DESEq2 documentation:
http://bioconductor.org/packages/release/bioc/vignettes/DESeq2/inst/doc/DESeq2.pdf

### D. Replicates evaluation

In order to analyze the similarity between the suggested replicates and explore the relations between the samples, heatmaps and PCA plots were generated. Principal component Analysis (PCA) generates a plot that span(s) the samples in 2D plane by their first two principal components.
